# Target cell-specific synaptic dynamics of excitatory to inhibitory neuron connections in supragranular layers of human neocortex

**DOI:** 10.1101/2020.10.16.343343

**Authors:** Mean-Hwan Kim, Cristina Radaelli, Elliot R. Thomsen, Deja Machen, Tom Chartrand, Nikolas L. Jorstad, Joseph T. Mahoney, Michael J. Taormina, Brian Long, Katherine Baker, Luke Campagnola, Tamara Casper, Michael Clark, Nick Dee, Florence D’Orazi, Clare Gamlin, Brian Kalmbach, Sara Kebede, Brian R. Lee, Lindsay Ng, Jessica Trinh, Charles Cobbs, Ryder P. Gwinn, C. Dirk Keene, Andrew L. Ko, Jeffrey G. Ojemann, Daniel L. Silbergeld, Staci A. Sorensen, Jim Berg, Kimberly Smith, Philip R. Nicovich, Tim Jarsky, Gabe Murphy, Hongkui Zeng, Jonathan T. Ting, Boaz P. Levi, Ed S. Lein

## Abstract

Rodent studies have demonstrated that synaptic dynamics from excitatory to inhibitory neuron types are often dependent on the target cell type. However, these target cell-specific properties have not been well investigated in human cortex, where there are major technical challenges in reliably identifying cell types. Here, we take advantage of newly developed methods for human neurosurgical tissue analysis with multiple patch-clamp recordings, *post-hoc* fluorescent *in situ* hybridization (FISH), and prospective GABAergic AAV-based labeling to investigate synaptic properties between pyramidal neurons and PVALB- vs. SST-positive interneurons. We find that there are robust molecular differences in synapse-associated genes between these neuron types, and that individual presynaptic pyramidal neurons evoke postsynaptic responses with heterogeneous synaptic dynamics in different postsynaptic cell types. Using molecular identification with FISH and classifiers based on transcriptomically identified PVALB neurons analyzed with Patch-seq methods, we find that PVALB neurons typically show depressing synaptic characteristics, whereas other interneuron types including SST-positive neurons show facilitating characteristics. Together, these data support the existence of target cell-specific synaptic properties in human cortex that are similar to rodent, thereby indicating evolutionary conservation of local circuit connectivity motifs from excitatory to inhibitory neurons and their synaptic dynamics.

## INTRODUCTION

Synaptic transmission is a fundamental means to convey information between neurons, and can be modulated by many factors including the intrinsic membrane properties of pre- and postsynaptic cell types, their connection probability, location of synapses, and synaptic short-term plasticity (STP) with timescales from milliseconds to minutes. Diverse forms of STP exist that involve differences in presynaptic release probability of neurotransmitters, calcium accumulation in presynaptic terminals, and retrograde signaling from postsynaptic dendrites with rapid timescales (Abbott and Regehr, 2004). Importantly, the properties of individual synapses from a given neuron are often determined by the identity of the postsynaptic neurons. Target cell-specific short-term synaptic dynamics from excitatory to inhibitory neuron connections have been identified in many brain regions including neocortex, cerebellum, and hippocampus (Blackman et al., 2013).

Rodent studies have begun to elucidate differential synaptic properties between specific neuron types, as well as their underlying postsynaptic molecular mechanisms. For example, specific postsynaptic molecules controlling presynaptic transmitter release have been identified, including N-cadherin and β-catenin (Vitureira et al., 2012), PSD-95-neuroligin (Futai et al., 2007), and Munc13-3 (Augustin et al., 2001). Excitatory to morphologically defined multipolar basket (or PVALB positive) cell synapses show a high initial release probability and synaptic depression. In contrast, excitatory to morphologically defined bi-tufted (or low threshold activated, SST positive) cell synapses show low initial release probabilities and synaptic facilitation (Reyes et al 1998; Koester & Johnston, 2005). This specialized short-term facilitation in SST interneurons is known to be mediated by Elfn1 (extracellular leucine rich repeat and fibronectin Type III domain containing 1) expression in postsynaptic dendritic shafts of SST cells (Sylwestrak & Ghosh, 2012; de Wit & Ghosh, 2016; Stachniak et al., 2019), but not in PVALB neurons. Elfn1 in postsynaptic SST neurons interacts with presynaptic metabotropic glutamate receptors (mGluRs) and kainite receptors in a layer-specific manner (Stachniak et al., 2019). Presynaptic mGluR localization opposed to SST neurons has also been reported in hippocampal pyramidal neurons (Scanziani et al., 1998; Shigemoto et al., 1996).

Addressing whether similar synaptic properties and molecular mechanisms are conserved in human cortex has been extremely challenging due to limitations in tissue access and available methods. Advances in single cell genomics have demonstrated a generally conserved cell type organization from mouse to human, but with many changes in cellular gene expression that suggest differences in cellular physiology, anatomy and connectivity (Hodge et al., 2019; Bakken et al., 2021). Recently, work from several research groups has shown that electrophysiological properties and local synaptic connectivity can be studied in acute human neocortical slices derived from surgical resections (Molnar et al., 2008; Jiang et al., 2012; Testa-Silva et al., 2014; Kalmbach et al., 2018; Beaulieu-Laroche et al., 2018; Boldog et al., 2018; Seeman et al., 2018; Peng et al., 2019; Planert et al., 2021; Campagnola, Seeman et al., 2022). These studies have demonstrated many conserved features, but a variety of human specializations compared to rodents, including faster recovery from synaptic depression (Testa-Silva et al., 2014) and greater numbers of functional release sites (Molnar et al., 2016).

The current study aimed to determine whether the target cell-dependent synaptic properties between excitatory pyramidal neurons and inhibitory PVALB^+^ vs. PVALB^−^ subclasses seen in rodent are conserved in human. We leveraged a number of technological advances to address this question, including 1) multiple patch-clamp (MPC) recordings to analyze intrinsic membrane properties and local synaptic connectivity and STP, 2) *post-hoc* multiplexed fluorescent *in situ* hybridization (mFISH) to reveal molecular properties of characterized neurons and synapses, 3) a novel human slice culture approach with cell class-specific adeno-associated virus (AAV) vectors to prospectively label GABAergic interneurons, and 4) a quantitative classifier to predict interneuron subclass identity based on a training set of human Patch-seq data with transcriptomically identified neurons (Lee et al., 2021). We find that STP in human cortex is target cell-specific. Excitatory to fast spiking (or PVALB positive) synapses show a high initial release probability and synaptic depression, whereas a subset of postsynaptic neurons with facilitating synapses were stained with SST by mFISH. Expression of *ELFN1* in human cortex is restricted to non-PVALB types similar to observations made in mouse, suggesting a conservation of molecular machinery mediating these target cell-specific synaptic properties.

## RESULTS

### Segregation of subclass level interneuron identities by synaptic membrane associated genes

Single-nucleus transcriptomic analyses from the neocortex of various species have identified a hierarchical classification of neuronal cell types that is conserved across cortical regions and species (Hodge et al., 2019; Bakken et al., 2021). This classification is consistent with a large literature describing stereotyped anatomy, physiology and connectivity, for instance for the major subclasses of cortical GABAergic neurons (e.g., PVALB, SST and VIP) (Paul et al., 2017; Huang and Paul, 2019). Importantly, transcriptomic analysis of the GABAergic subclasses in mouse cortex shows they are well differentiated from one another by genes involved in synaptic communication (Paul et al., 2017; Huang and Paul, 2019; Smith et al., 2019), suggesting a molecular substrate for their distinctive features of functional synaptic communication. Here we used human and mouse single cell/nucleus RNA-seq data to compare gene expression between GABAergic SST and PVALB subclasses and between species. We identified 72 PVALB- and 75 SST-enriched genes whose expression patterns were conserved in both human medial temporal gyrus (MTG) and mouse primary visual cortex (VISp) (Tasic et al., 2018; Hodge et al., 2019; **Figure 1a**). These patterns were similar in primary motor cortex (M1 in human, MOp in mouse), consistent with reports of similar transcriptomic GABAergic neuron type properties across mouse cortical areas (Tasic et al., 2018). For example, the *ELFN1* gene, described to mediate selective short-term facilitation in SST interneurons (Sylwestrak & Ghosh, 2012; de Wit & Ghosh, 2016; Stachniak et al., 2019), is enriched in GABAergic interneurons compared to excitatory neurons, and in SST and all other GABAergic subclasses except PVALB interneurons, and this pattern is conserved across both species and cortical areas.

**Figure 1.**
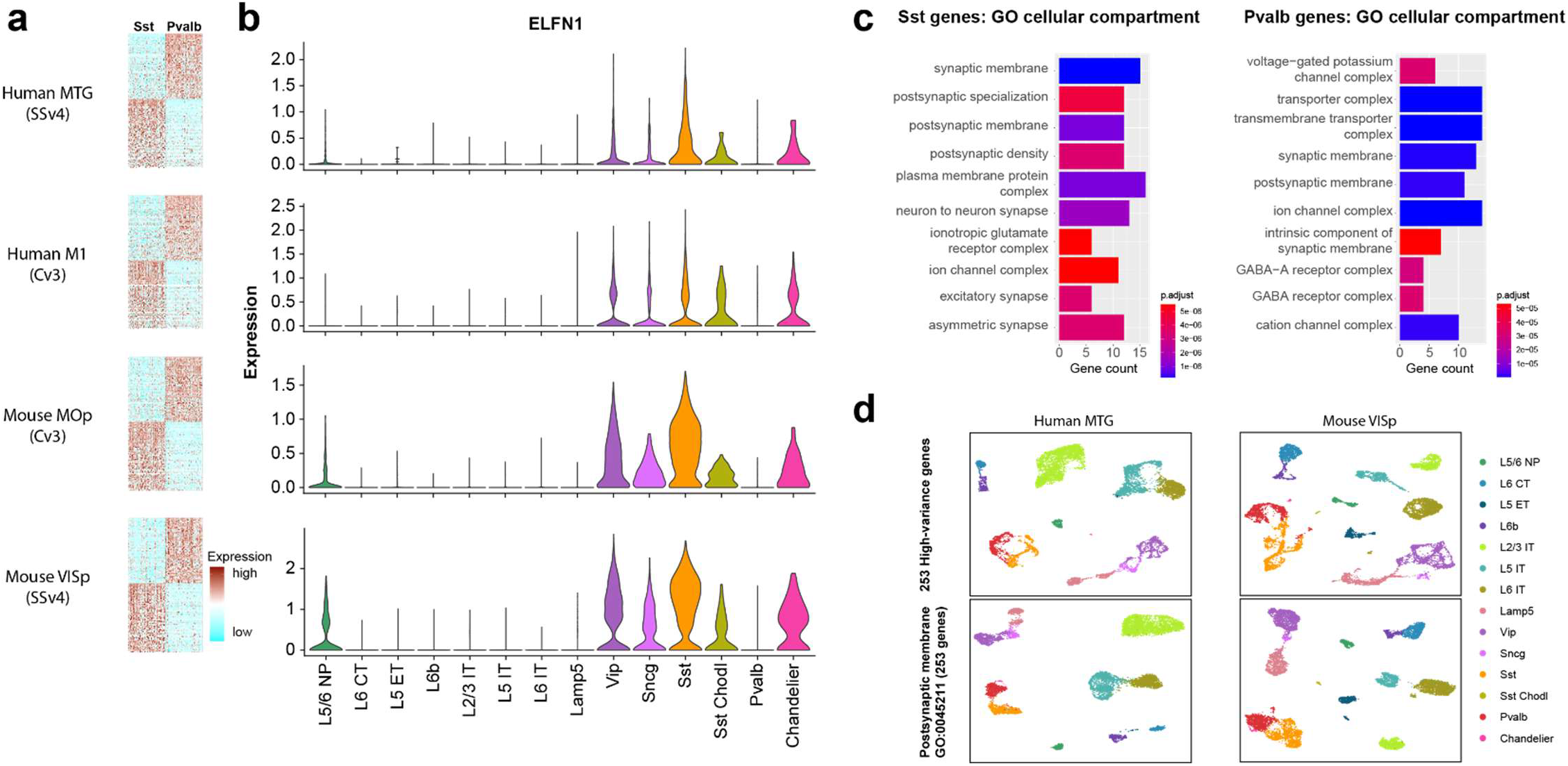
Single-nucleus transcriptomic differences between PVALB and SST types in human and mouse cortex. **a.** Heatmaps showing scaled log2 normalized expression of 147 differentially expressed genes (DEGs) that distinguish PVALB and SST types in both human MTG and mouse VISp. These genes show similar specificity in human M1 and mouse MOp, indicating conserved patterning across cortical areas. Heatmaps show 100 randomly sampled nuclei from each type. SSv4 indicates SMARTseq V4 chemistry, and Cv3 indicates 10x Chromium V3 chemistry. **b**. Violin plots showing neuronal subclass expression levels of *ELFN1*, illustrating selective expression in non-PVALB inhibitory neuron subclasses. **c.** Gene ontology analysis for cellular compartment using conserved SST or PVALB DEGs. Top 10 enriched categories are involved in synaptic structure and function. **d**. Similar cellular architecture shown with UMAPs constructed only using postsynaptic membrane GO: 0045211 term genes for human and mouse neurons (top panels), compared to UMAPs constructed using a comparably sized set of 253 high-variance genes (bottom panels).

Gene ontology (GO) analysis for cellular compartment was performed on the 72 PVALB and 75 SST enriched genes independently, identifying numerous significantly enriched categories (**Figure 1c**, top 10 categories by p-value). The top terms were strongly enriched for synapse related categories, with postsynaptic membrane term GO:0045211 being enriched in both PVALB and SST neurons, suggesting differences in their synaptic connectivity and functional properties. The typical approach to clustering and representing transcriptomic data is to cluster based on unbiased sets of high variance genes, as shown in the upper panels of **Figure 1d**. Remarkably, clustering only using genes in the postsynaptic membrane term GO:0045211 produces a very similar result that clearly differentiates all of the glutamatergic and GABAergic subclasses (**Figure 1d**, lower panel). These analyses indicate that each neuronal subclass expresses unique combinations of conserved synaptic membrane associated genes that underlie their unique development of synaptic formation and functional properties (Blackman et al., 2013). On the other hand, there are also many genes that differentiate PVALB and SST subclasses in mouse and in human, but that are not conserved between species. These include genes that are associated with neurotransmitter signaling pathways and synapse function (**Figure 1 – Figure supplement 1**), suggesting that there are also species specializations in GABAergic subclass-specific synaptic function.

### Local synaptic connectivity and intrinsic membrane properties in acute and virally transduced neurons from human *ex vivo* cultured cortical slices

Next, we investigated local synaptic dynamics from excitatory to inhibitory neurons in supragranular layer of human neocortex using multiple whole-cell patch-clamp (MPC) recordings. In this study, neocortical tissues from 31 donors were used for data collection, derived from neurosurgical resections to treat intractable epilepsy (*n* = 15 cases, *n* = 59 connected pairs) or remove deep brain tumors (*n* = 16 cases, *n* = 30 connected pairs). These tissues have been shown to exhibit minimal pathology, are distal to the epileptic focus or tumor, and have been used extensively to characterize cellular physiology and anatomy previously (Berg et al., 2021). Donors included males and females across adult ages, and tissues from left and right hemispheres and temporal and frontal cortices (**Figure 2 – Figure supplement 1**).

Two main experimental approaches were applied, including an acute brain slice preparation and an organotypic brain slice culture preparation (**Figure 2**). Notably, both applications were typically performed on the same surgical cases, since multiple slices could be generated from these resections whose average volume was 1.39 ± 0.57 cm^3^ (mean ± standard error of mean (s.e.m); averaged over *n* = 12 cases). Acute experiments were performed within 12 hours following surgical resection, whereas slice culture experiments were performed between 2.5-9 days *in vitro* (DIV; **Figure 2**). In acute slice preparations, neurons were targeted based on somatic shape as visualized by oblique illumination. In slice culture experiments, AAV vectors were used to drive fluorescent reporters under the control of cell class-selective regulatory elements to facilitate targeting labeled neurons for MPC recordings. After MPC electrophysiology experiments two strategies were used to identify GABAergic subclass identity, including direct mFISH analysis with subclass markers, and a computational classifier based on Patch-seq analysis using similar human slice preparations.

**Figure 2.**
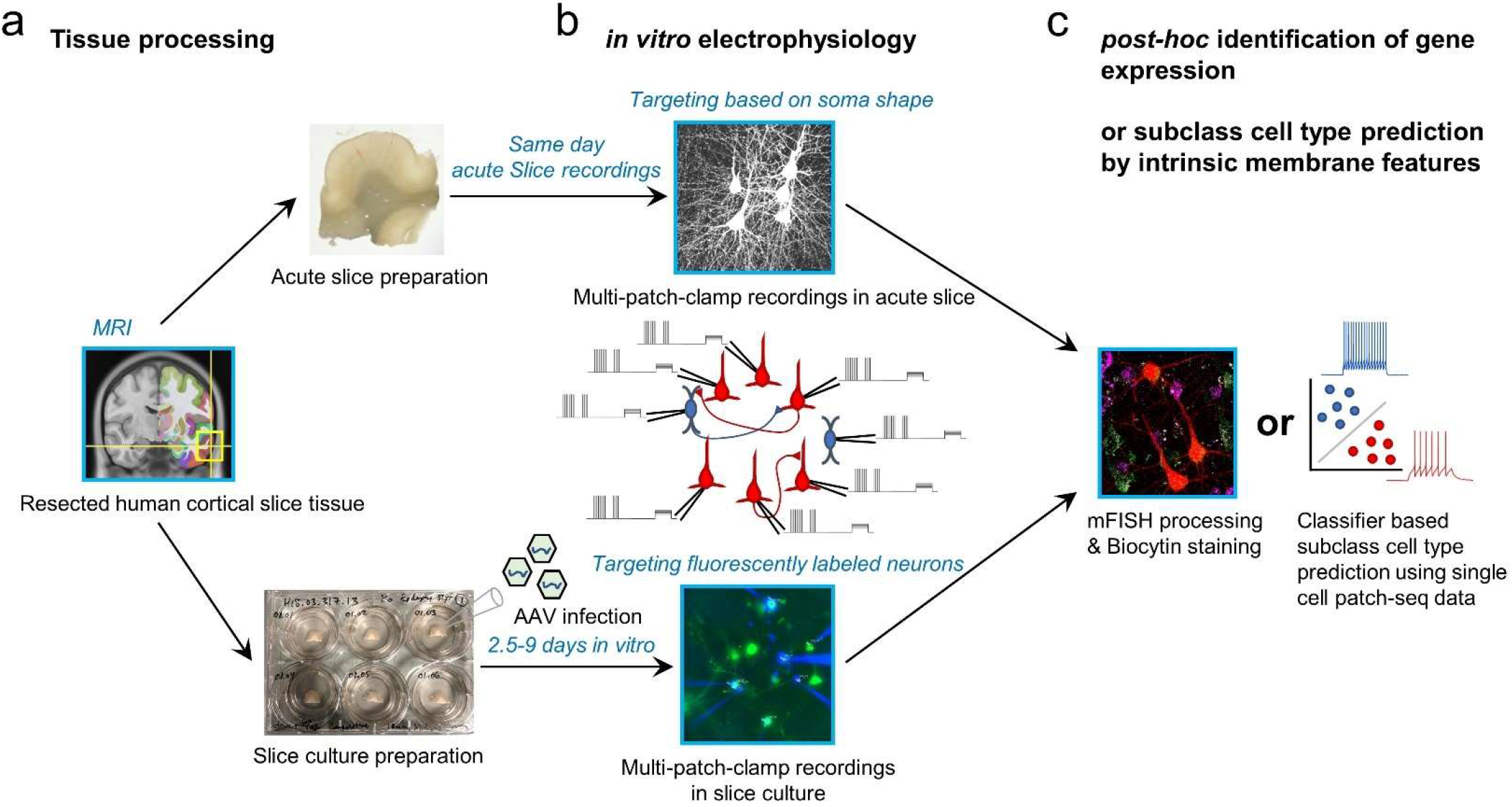
Schematic of experimental workflow. **a**, Human neocortical tissue from neurosurgical resections enter either acute slice preparations within 45 min following scalpel excision from the patient (upper) or organotypic slice culture preparation with viral transduction (lower). **b**, Up to eight simultaneous patch-clamp recordings are performed on either acute slices (upper) or slice culture after 2.5 to 9 days *in vitro* (lower). Targeting of neurons is either carried out by visually identifying cell bodies using an upright microscope with oblique illumination (upper) or by targeting neurons expressing fluorescent reporters following viral infection (lower). **c**, To identify subclass cell types in connectivity assayed neurons, we applied multiplexed fluorescence *in situ* hybridization (mFISH) on fixed slices to identify marker gene expression, as well as a machine learning classifier with cellular intrinsic membrane properties measured after connectivity assays.

Connectivity assays with MPC recordings were performed by targeting cell bodies located between 50 and 120 µm below the surface of the slice to minimize truncation of dendrites and other superficial damage that occurs during slice preparation (**Figure 3a-g**; Seeman et al., 2018; Campagnola, Seeman et al., 2022). To look at synaptic connections between excitatory pyramidal and inhibitory interneuron in acute experiments, we simultaneously targeted cells with pyramidal shape in addition to small round, putative interneurons. For example, **Figure 3** shows an MPC experiment that successfully targeted 4 pyramidal neurons (shown with reconstruction of biocytin staining; **Fig. 3i**) and one interneuron. Connectivity was observed from excitatory to inhibitory for the interneuron, which displayed fast spiking characteristics (cell 4), with strong excitatory postsynaptic potential (EPSP) responses that rapidly depress (e.g., cell 1 to cell 4 and cell 2 to cell 4). In contrast, the pyramidal to pyramidal responses were small with weakly depressing characteristics (e.g., cell 3 to cell 5; **Figure 3b,c**). The intrinsic membrane features (**Figure 3d-g**) and morphology (**Figure 3h,i**) of this interneuron were consistent with the identity of a PVALB cell type (Reyes et al., 1998).

**Figure 3.**
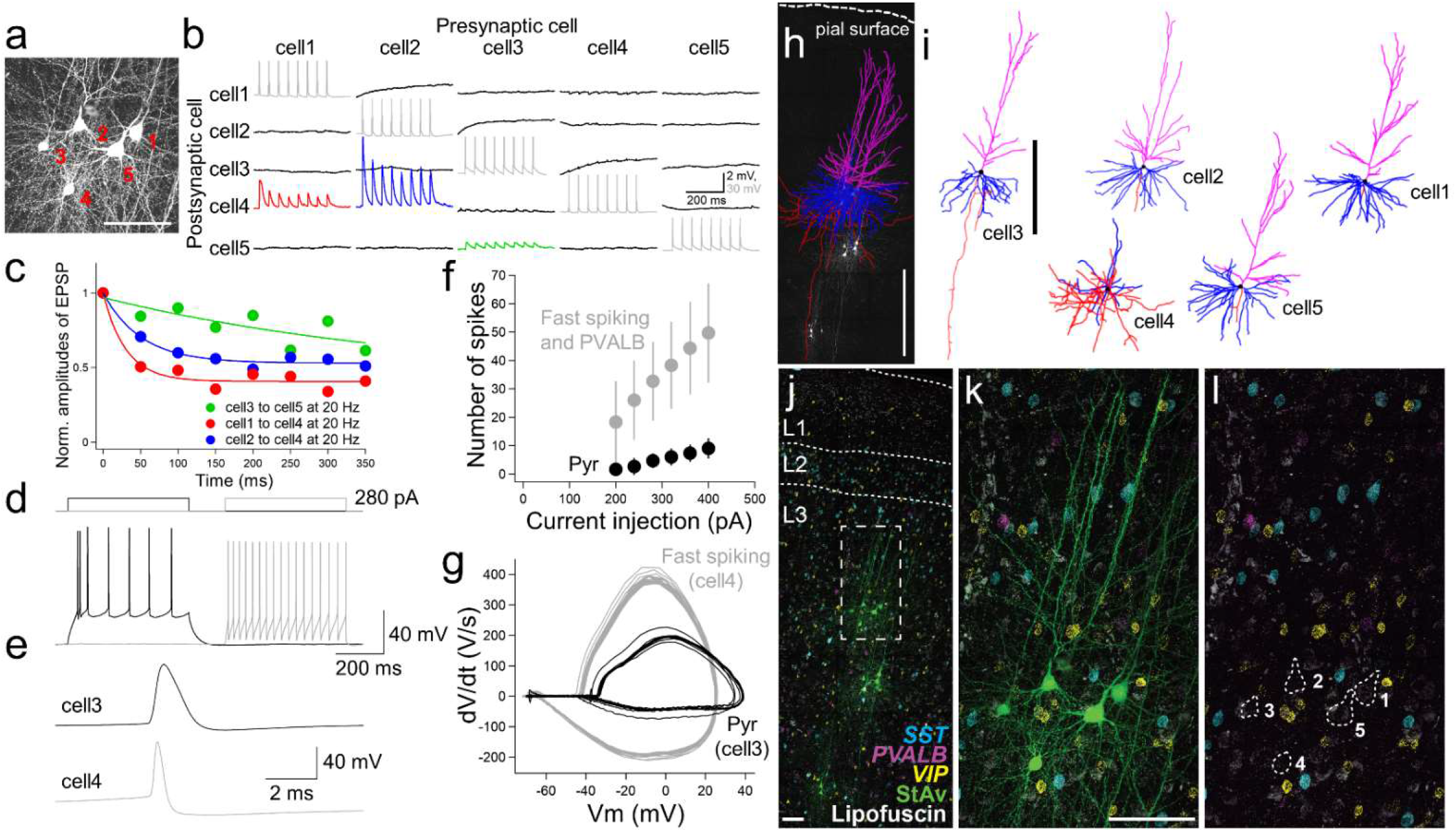
Quadruple modality data in acute *ex vivo* human neocortex. Example experiment using acute slice preparation with five cells simultaneously patched. **a**, Maximum intensity projection montage confocal image of biocytin/streptavidin labeling. Scale bar, 50 µm. **b**, Corresponding membrane voltage traces from connectivity assay. Presynaptic action potentials (gray) in individual neurons (cell1 to cell5) were sequentially generated by 8 brief current pulses at 20 Hz while simultaneously recording the postsynaptic membrane voltage in non-stimulated neurons in current-clamp mode (black). Traces averaged over 10 repetitive 8 pulse stimulations. This probing uncovered a strong and adapting excitatory synaptic connection from cell 2 to cell 4 (blue trace) and cell1 to cell4 (red trace) compared to the synaptic connection from cell3 to cell5 (green). **c**, Summary plot of short-term synaptic dynamics with presynaptic 20 Hz stimulation (8 pulses at 20 Hz) in connected pairs as in **b**. Amplitude normalized to size of initial EPSP. **d**, Example traces of action potential generation by step current injection in regular spiking (cell3, black) and fast spiking neurons (cell4, gray). The same amount of current injection (280 pA) was applied to cell3 and cell4. **e**, Spike shape comparison between regular and fast spiking neurons detected in the connectivity assay shown in **b**. **f**, Frequency-current curve of pyramidal neuron (Pyr; mean ± standard deviation, *n* = 3), and fast spiking neuron (panel **k**, cell4) and PVALB positive neurons (including upper 2 cells shown in panel **g** of **Figure 5 – Figure supplement 1**) (mean ± standard deviation, *n* = 3). **g**, Phase plot (dV/dt vs V) analysis based on responses shown in **d**. **h**, Morphological reconstruction of the 5 recorded neurons shown in **a**. Scale bar, 500 µm. **i**, Reconstruction of individual neurons. Scale bar, 500 µm. Blue, magenta, and red indicate basal dendrites, apical dendrites, and axons in pyramidal neurons (cell 1,2,3,5). For the interneuron (cell4), blue and red indicate dendritic and axonal structures, respectively. **j,** Fluorescence montage of cells imaged in **a, j-l** stained by mFISH for inhibitory neuron subclass markers (*PVALB*, *SST*, and *VIP*) and biocytin. MPC recordings were performed on three separate cell clusters in this slice (**j**). Note, substantial lipofuscin is observed in this slice. White box in **j** is shown at higher magnification for mFISH and biocytin (**k**), or mFISH only (**l**).

mFISH was used to confirm *PVALB* mRNA expression in this cell. One challenge to this approach is that human brain tissue often exhibits dense lipofuscin around some somatic structures (**Figure 3j-l, Figure 5 – Figure supplement 1e-g**), and persists after tissue clearing with 8% SDS and throughout the mFISH staining procedure. However, it was possible to distinguish the distribution of amplified mRNA fluorescent dots from lipofuscin autofluorescence by imaging across multiple channels, as lipofuscin produced fluorescent in all channels (e.g., **Figure 5 – Figure supplement 1e-g**). In this case PVALB labeling was observed in this cell, but the staining was very weak. We observed this with several patched PVALB*^+^* cells, where *PVALB* mRNA abundance was at lower levels than adjacent unpatched PVALB*^+^*cells (**Figure 5 – Figure supplement 1e-g**). Whether this reflects real differences in mRNA abundance between cells, or dilution or leakage of mRNA during MPC recording is unclear.

To efficiently target GABAergic interneurons for MPC recordings, we also performed rapid viral genetic labeling of cortical GABAergic interneurons in human organotypic slice cultures (see **Methods**; Ting et al., 2018; Mich et al., 2021). We used an adeno-associated virus (AAV) that drives SYFP2 reporter expression under the control of an optimized version of a previously described forebrain GABAergic neuron enhancer (Stuhmer et al., 2002; Dimidschstein et al., 2016). This AAV-DLX2.0-SYFP2 virus was directly applied to the slice surface at a concentration of 1-5e^10^ vg/slice. Fast reporter expression allowed us to execute physiology experiments after 2.5 days *in vitro* (DIV) onward after viral administration. We performed targeted MPC recordings of labeled neurons in addition to pyramidal shape neurons in human cortical slices (**Figure 5 – Figure supplement 1**). We noted some differences between MPC recordings in viral labeled slice culture and acute slice preparation. First, giga-ohm seals were more readily obtained between patch pipette and cell membrane in neurons from *ex vivo* cultured slices compared to acute slices. Second, the somatic structure of unlabeled neurons was more difficult to resolve in slice culture with minimal positive pressure on the patch pipette, making patching unlabeled neurons more challenging. Nonetheless, the ability to exclusively target genetically labeled GABAergic neuron subclasses in the human neocortex greatly improved throughput and efficiency of targeted recording experiments.

In human organotypic slice cultures obtained in this study (4-9 DIV), we generated reliable action potentials by a brief current injection on the patched presynaptic soma, and performed subsequent synaptic connectivity assays in both excitatory to excitatory and excitatory to inhibitory connected pairs. In acute slices, unitary presynaptic action potentials can evoke postsynaptic spikes in human basket cells (Molar and Tamas 2008, Szegedi et al., 2017) and in non-human primate acute cortical slice recordings. In human slice culture, we often observed a similar result from excitatory to inhibitory connections, even at −70 mV holding potential (**Figure 4 – Figure supplement 1**), suggesting that these synaptic properties are preserved.

### Diverse synaptic dynamics from excitatory to inhibitory neuron connections in supragranular layer of human cortex

To analyze excitatory postsynaptic potential (EPSP) dynamics, we stimulated presynaptic neurons with spike trains of 8 pulses at 20 and 50 Hz (see **Methods**, **Figure 4a**, Seeman et al., 2018; Campagnola, Seeman et al., 2022). Recovery from synaptic depression was measured by probing with an additional four pulses after variable inter-spike intervals (62.5 ms, 125 ms, 250 ms, 500 ms, 1 s, 2 s and 4 s) following induction by the 8 pulses spike train at 50 Hz. In our MPC recordings, at least three cells were patched simultaneously, and we either simultaneously patched two presynaptic pyramidal neurons and one connected postsynaptic interneuron (**Figure 4b,c**), or one pyramidal neuron and two connected postsynaptic interneurons (**Fig. 4d**). As shown in **Figure 4b** and **4c**, when two pyramidal neurons were connected to the same postsynaptic inhibitory neuron, they typically showed similar kinetics of short-term synaptic plasticity that was either depressing or facilitating depending on the postsynaptic neuron. Similarly, when one presynaptic pyramidal neuron was connected to 2 interneurons, the short-term synaptic plasticity was often different for the two interneurons (**Fig. 4d**). Both of these results indicate that postsynaptic cell identity is a determinant of short-term synaptic dynamics (i.e., target cell-specific) in excitatory to inhibitory neuron connections in human cortex (**Figure 4b-d**; Reyes et al 1998; Koester & Johnston, 2005). These target-dependent synaptic properties were observed in both acute and slice culture preparations.

**Figure 4.**
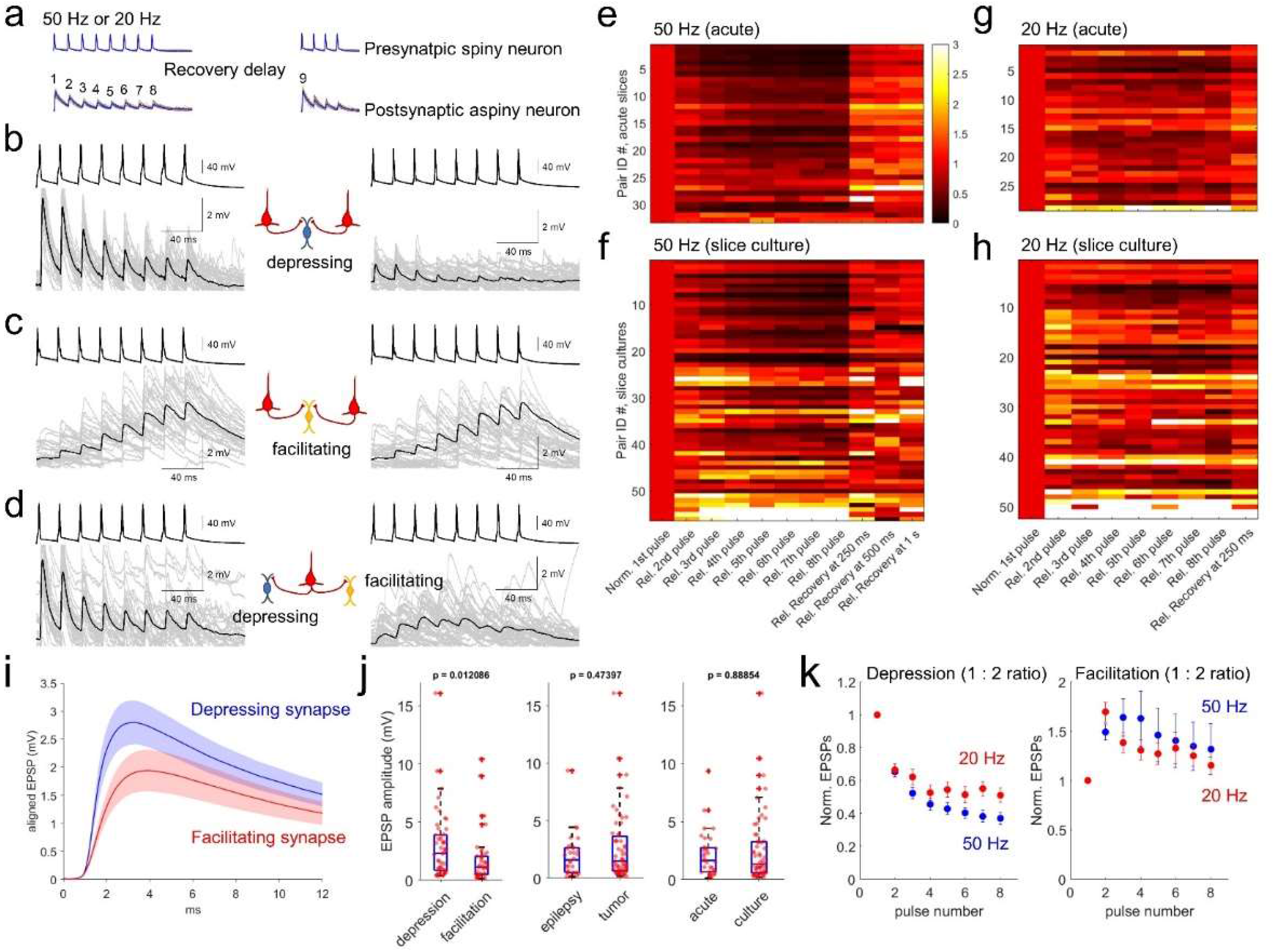
Survey of synaptic dynamics measured from supragranular pyramidal neurons to neighboring interneurons in human cortex. **a,** Stimulation protocol of connectivity assays. Eight trained presynaptic spikes were generated with two main stimulus frequencies (20 and 50 Hz). Fixed 250 ms recovery delay was used for 20 Hz stimulation and a range of recovery delays (from 62.5, 125, 250, 500 ms, 1 s, 2, and 4 s) were interposed between the eight induction pulses and four recovery pulses. **b-d,** Target cell-specific synaptic dynamics from pyramidal to interneuron connections. Two pyramidal neurons were connected to one interneuron and their synaptic dynamics were similar, i.e., both were depressing (**b**), or both were facilitating (**c**). However, example in **d** shows one pyramidal neuron that was connected to two interneurons showing either depression (left panel) or facilitation (right panel). Averaged EPSP responses (blue, thick line) on top of individual responses traces (multiple colors) are displayed in each connected pair. **e-h,** Initial EPSP sizes were normalized in connected pairs, and their relative synaptic dynamics according to presynaptic train stimulation (8 pulses) are displayed by heatmap. Heatmap rows sorted based on the size of EPSP, from largest (top row) to smallest (bottom row). Initial recovery pulse (denoted as 9 in panel **a**) responses out of 4 recovery pulses are displayed at 250 ms, 500 ms and 1 second at 50 Hz (**e,f**) and fixed recovery interval at 250 ms at 20 Hz (**g,h**) as example responses. Synaptic connections found in acute slices (*n* = 33 at 50 Hz in panel **e**; *n* = 29 at 20 Hz in panel **g**) and slice culture (*n* = 56 at 50 Hz in panel **f**; *n* = 52 at 20 Hz in panel **h**). **i**, Initial EPSP responses from 50 Hz stimulation were aligned from response onset and averaged. 1:2 ratio was determinant for classifying depression and facilitation at 50 Hz presynaptic stimulation. Aligned average EPSP kinetics are shown (depressing synapses, *n* = 50; facilitating synapses, *n* = 39). Displayed data indicate mean (blue, red) ± s.e.m (shaded regions with light colors). **j**, Amplitudes of EPSP responses (i.e., averaged first EPSP responses at 50 Hz stimulation in connected synapses) were compared between depressing (*n* = 50) and facilitating (*n* = 39) synapses defined by their 1:2 ratio at 50 Hz stimulation. EPSP amplitudes were compared from their tissue origins (*n* = 30, epilepsy; *n* = 59, tumor). EPSP amplitudes were also compared from their tissue preparation types (*n* = 33, acute slice; *n* = 56, slice culture). P-values are from Wilcoxon rank sum test. **k**, Kinetics of synaptic dynamics (1 to 8 pulses, normalized to first response) were compared at different frequencies (i.e., 50 Hz and 20 Hz presynaptic stimulation). Depression and facilitation synapses were defined based on 1:2 ratio. Kinetics of dynamics from depressing synapses are displayed (mean ± s.e.m; *n* = 50, depression at 50 Hz, blue; *n* = 29, depression at 20 Hz, red; left). Similarly, kinetics of dynamics from facilitating synapses are displayed (*n* = 39, facilitation at 50 Hz; *n* = 50, facilitation at 20 Hz; right).

Synaptic dynamics of connected excitatory to inhibitory neuron pairs were analyzed from both acute (*n* = 33 at 50 Hz; *n* = 29 at 20 Hz stimulation protocol) and slice culture preparations (*n* = 56 at 50 Hz; *n* = 52 at 20 Hz stimulation protocol). To quantify synaptic dynamics, initial EPSP amplitudes in each pair of excitatory to inhibitory neuron connections were normalized and displayed as heatmaps (**Figure 4e-h**). Rates of postsynaptic facilitation and depression are usually presynaptic stimulus frequency dependent (Beierlein et al., 2003). Here, 50 Hz stimulation protocol showed stronger depression with bigger EPSP responses (upper heatmap in both acute and slice culture data, **Figure 4e,f**) compared to the 20 Hz stimulation protocol (**Figure 4g,h**). These results suggest that presynaptic vesicle pools could be more quickly replenished with lower frequency presynaptic stimulation (Waters and Smith, 2002). Connected pairs with bigger EPSPs (upper heat map at 50 Hz; **Figure 4e,f**) tended to have depressing synapses, whereas connected pairs with smaller EPSPs (lower heat map at 50 Hz; **Figure 4e,f**) tended to have facilitating synapses, suggesting that the large and small EPSP synapses may represent different inhibitory neuron types.

We did observe some differences in synaptic properties between AAV-labeled GABAergic interneurons in slice culture and putative interneurons in acute slices targeted based on homogeneous small round soma shape under the microscope. Initial EPSP amplitudes were not significantly different between slice cultures and acute slices (**Figure 4j,** right panel). In contrast, we observed that normalized synaptic dynamics were significantly different between acute and slice culture preparations (see 50 Hz, left panel of **Figure 4 – Figure supplement 4d**). More facilitating synapses were detected in slice cultures than in acute slices. Based on the train-induced STP (1 : 6-8 ratio), about 30% of recordings (*n* = 17) in slice cultures (total *n* = 56) showed facilitation, compared to only 12% of recordings (*n* = 4) in acute slices (total *n* = 33) (**Figure 4f,h**). This difference could either reflect an acute vs. slice culture difference, or more likely a selection bias for interneuron subtype sampling between slice preparation methods as discussed below.

To further explore the potential relationship between interneuron subtypes and synaptic properties, we first defined synapses as facilitating or depressing based on the ratio of EPSP amplitude between the 1^st^ and 2^nd^ pulse (1:2 ratio) (**Figure 4k** right panel, **Figure 4 – Figure supplement 4a,b** right panel, Beierlein et al., 2003). The depressing synapses had significantly higher EPSP amplitudes than facilitating synapses (**Fig. 4j**, left panel), and this difference was not accounted for by disease indication or slice preparation method. Specifically, we did not observe significantly different dynamic responses related to these variables (i.e., normalized responses from first to 8^th^ pulses at both 20 Hz and 50 Hz; **Figure 4 – Figure supplement 4c**). EPSP amplitudes and their recovery responses (i.e., 9^th^ pulse response) at various time intervals were not significantly different when we compared based on their tissue origins (i.e., epilepsy vs tumor case; **Figure 4 – Figure supplement 3b**) and preparation (i.e., acute vs slice culture; **Figure 4 – Figure supplement 3c**).

These observed differences in pyramidal to interneuron synaptic properties could relate to previously described differences in pyramidal neuron to PVALB-positive interneuron (depressing) and SST-positive (facilitating) interneurons (Reyes et al 1998; Koester & Johnston, 2005). In mouse V1, EPSP rise time and EPSP decay tau is shorter in pyramidal to PVALB neurons compared to pyramidal to SST neurons in mouse V1 (Campagnola, Seeman et al., 2022). In human cortex, we also see a trend towards differential kinetics for EPSP rise time and decay between depressing and facilitating synapses as shown in averaged responses (**Figure 4i**), although those responses were not statistically different (Wilcoxon rank sum test, **Figure 4 – Figure supplement 2**). Frequency dependent lateral inhibition between neighboring pyramidal neurons through facilitating Martinotti cells has been reported in both rodents (Silberberg and Markram, 2007; Berger et al., 2009) and human (Obermayer et al., 2018). We saw a slight trend towards activity dependent facilitation in facilitating synapses defined by 1:8 ratio (**Figure 4 – Figure supplement 4a** right panel), but this was not clear in other analyses (i.e., 1:2 pulse ratio, or 1:6-8 pulse ratio; **Figure 4k** right panel**, Figure 4 – Figure supplement 4b** right panel). Therefore, there are trends to indicate that high EPSP amplitude, depressing synapses in human cortex may represent excitatory synapses onto PVALB-positive neurons, and low EPSP amplitude, facilitating synapses may represent excitatory synapses onto Somatostatin-positive neurons.

### Subclass level cell type identification of postsynaptic interneurons in MPC by *post-hoc* HCR staining

We used hybridization chain reaction (HCR) mFISH to identify postsynaptic interneuron subclass identities following human multipatch experiments. This method was used because it penetrates tissue efficiently (Choi et al., 2010), allows strong signal amplification, has high signal-to-noise with background-reducing probe design (Choi et al., 2018), and allows multiple rounds of stripping and re-probing (**Figure 5 – Figure supplements 1,2,3**. Following MPC recordings, slices were fixed, passively cleared, and stained by mFISH using HCR kit version 3.0 (Shah et al., 2016; Choi et al., 2018). Messenger RNA from excitatory (*SLC17A7*) and inhibitory (*GAD1*) marker genes were easily resolved in both patched (biocytin/streptavidin, StAv) and neighboring non-patched neurons (**Figure 5 – Figure supplement 1a-d**). As expected, *SLC17A7* and *GAD1* expression was mutually exclusive in excitatory and inhibitory neurons, respectively, and only GAD1^+^ cells were found in layer 1. In general, *SLC17A7* and *GAD1* mRNA staining was comparable between patched and neighboring non-patched neurons after long whole-cell recordings (around 30-75 min; **Figure 5 – Figure supplement 1b,c,i,j**). However, there was trend for lower signal detection for GABAergic markers PVALB and SST in patched versus non-patched cells (**Figure 5 – Figure supplement 3**), indicating that there may be some dialysis or degradation of mRNA from the cell during recording but that it affects detection of different genes differentially. We were also able to resolve *SLC17A7* and *GAD1* mRNA staining through the depth of the slice, and did not observe significant changes of averaged fluorescent intensities in individual neurons by depth (**Figure 5 – Figure supplement 1j,k**). We also did not observe any noticeable difference in mRNA staining intensity between patched and neighboring non-patched neurons in slice culture (**Figure 5 – Figure supplement 1, 3a**). The ability to stain across multiple rounds allowed probing for an increased number of genes, and re-probing for genes that produced low signal from the first round such as *VIP* (**Figure 5 – Figure supplement 2c-d** cell3).

As shown in **Figure 5 – Figure supplement 2a,b**, the use cell class specific AAV vectors facilitates efficient prospective cell class labeling and subsequent identification of cortical interneuron subclasses that are difficult to reliably target in acute brain slice preparations. All ten patched cells were GABAergic, and multiple VIP, SST, and PVALB-positive neurons were patched from two sets of MPC recording attempts (site 1 and 2 in **Figure 5 – Figure supplement 2a,b**).

Using *post-hoc* HCR in connectivity assays with MPC recordings in thick tissue human acute and slice culture preparation, we were able to identify postsynaptic GABAergic subclass identity in connected pairs with presynaptic excitatory neurons (**Figure 5**). Initial EPSP amplitudes in the pairs with postsynaptic PVALB positive neurons were significantly larger compared to those of postsynaptic SST positive neurons (EPSP amplitudes, mean ± s.e.m, 4.7393 ± 1.3975 mV with PVALB positive postsynaptic neuron, 0.8412 ± 0.3147 mV with SST positive postsynaptic neuron, P = 0.0159, Wilcoxon rank sum test). Whereas synaptic dynamics with PVALB positive neurons shows strong depression, synaptic dynamics with SST positive neurons shows facilitation and their normalized trained responses are significantly different (i.e., 2-8 pulse responses in **Figure 5f**; *, P < 0.05, Wilcoxon rank sum test, *n* = 5 for PVALB and *n* = 6 for SST).

**Figure 5.**
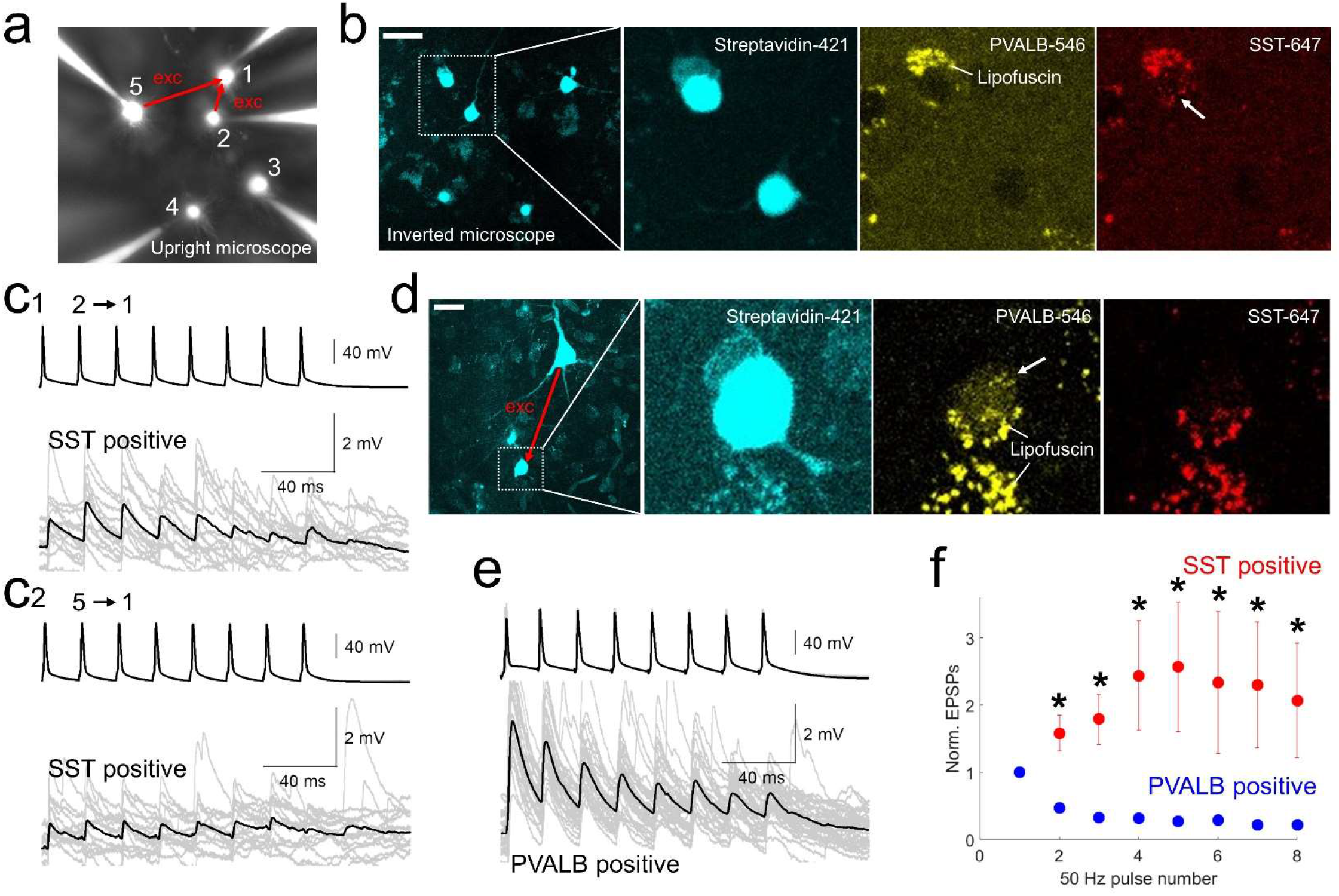
*post-hoc* HCR (mFISH) staining confirmed differentiated synaptic dynamics from excitatory pyramidal neurons to PVALB (depressing) and SST (facilitating) positive interneurons. **a-b**. MPC recordings were performed underneath upright microscope (cascade blue, fluorescent dye included with biocytin on the patch pipettes, **a**), and biocytin filled patched cells, stained with streptavidin (left two panels, **b**) and their HCR staining were identified on the inverted microscope. Example of SST-stained neuron (cell1) in connectivity assay from slice culture. Note that lipofuscin signal was seen in both PVALB (546 nm) and SST (647 nm) channels, but HCR signals were shown up separately in one or the other channel. **c1-2**, Corresponding synaptic dynamics at 50 Hz stimulation. In this example, two pyramidal neurons (cell2 and cell5) were connected to a SST-positive neuron (cell1). **d**, Example of PVALB-stained neuron in connectivity assay from acute slice (**d**). **e**, Corresponding synaptic dynamics at 50 Hz presynaptic stimulation is displayed. Averaged EPSP responses (black, thick line) on top of individual responses traces (gray) are displayed for each connected pair in (**c1, c2**) and (**e**). **f**. Summary plot of normalized synaptic dynamics from PVALB positive (blue; *n* = 6) and SST positive (red; *n* = 6) neurons. Displayed data indicate mean ± s.e.m (error bars; *, p < 0.05, Wilcoxon rank sum test).

### Cell subclass identification of postsynaptic interneurons by machine learning classifier

Since *post-hoc* HCR on MPC experiments is such a low-throughput method, we took advantage of a larger existing human single cell Patch-seq datasets to develop a quantitative classifier to predict interneuron subclass identity on our larger MPC dataset. This reference dataset comprised a set of Patch-seq experiments in slice culture that targeted AAV-DLX2.0-SYFP2 labeled neurons (Berg et al., 2021, Lee et al., 2021; see **Methods**), from which the cells were robustly defined based on transcriptomic analysis following electrophysiological characterization, nucleus extraction and RNA sequencing. Such a classifier strategy was possible because intrinsic membrane properties of each cell were measured in our connectivity assays with MPC recordings, including subthreshold step hyperpolarization and depolarization from −70 mV holding potential and suprathreshold step depolarization (**Figure 3d,e,f,g**, and **Figure 6c**). There were a few noticeable differences between MPC recordings and single cell Patch-seq experiments such as composition of internal solution (e.g., RNAase inhibitor, EGTA concentration, etc.) and external aCSF solution (e.g., synaptic blockers and Ca^2+^ concentration), and fewer stimulus protocols of intrinsic properties in MPC recordings (in addition, maintaining iso-potential around −70 mV in MPC recordings, rather than resting potential in Patch-seq recordings) (see **Methods** for the details). To control for these methodology-based differences, the two datasets were pre-aligned using supervised feature selection prior to classifier training and application (see **Methods**).

**Figure 6.**
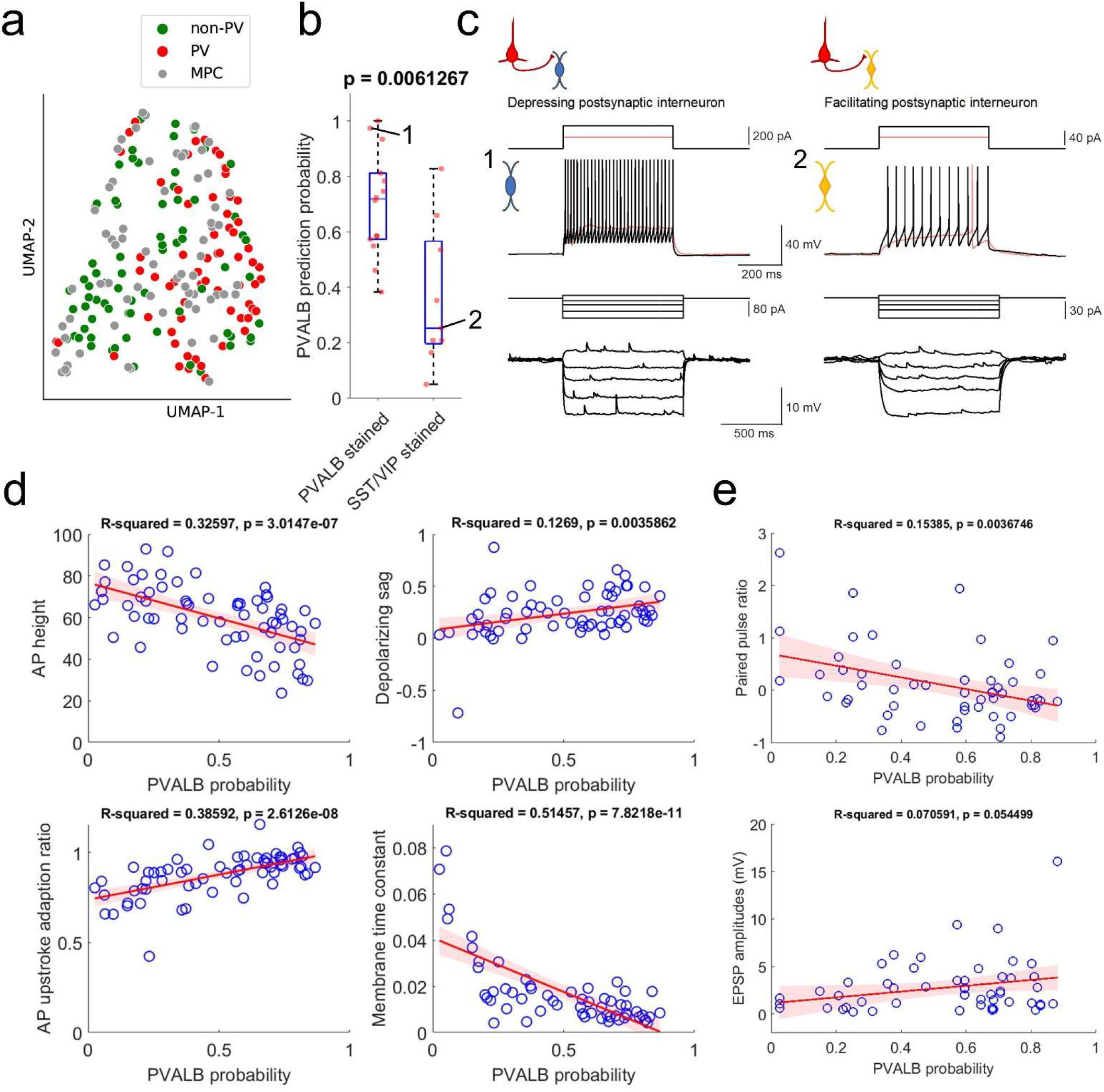
Prediction of PVALB and non-PVALB subclass cell type identities in postsynaptic interneurons by intrinsic membrane properties-based classifier using human Patch-seq data obtained from AAV virally labeled slice cultures. **a**, UMAP visualization of PVALB and non-PVALB cell types based on their intrinsic membrane properties using human single cell Patch-seq data and alignment with postsynaptic cell intrinsic properties from connectivity assay (MPC). **b**, Correlation between PVALB probability predicted by intrinsic properties-based classifier and their subclass identity based on HCR staining (PVALB stained, *n* = 14; SST/VIP stained, *n* = 9). Classification based on PVALB and non-PVALB cell prediction. P value from Wilcoxon rank sum test. **c**, Example traces of intrinsic membrane properties from postsynaptic cells, showing synaptic depression (cell1, PVALB positive) and facilitation (cell2, SST positive). **d**, Examples of identified dominant features to segregate PVALB and non-PVALB interneurons. **e**, Correlation between PVALB probability and their paired pulse (1:2 ratio, upper) or EPSP amplitudes (lower) at 50 Hz presynaptic stimulation protocol. Regression line (red) with fitting confidence bounds (shaded region) in **d,e**.

Using only these electrophysiological features, it was possible to differentiate between PVALB-positive GABAergic interneurons and other, non PVALB-positive interneurons, as illustrated in the UMAP in **Figure 6a**. The MPC recordings were intermingled with the Patch-seq neurons, indicating overlapping properties and coverage of both cell groups across the two datasets. Importantly, quantitative predictions for the PVALB identity of postsynaptic interneurons from MPC recordings using a classifier trained on these intrinsic features matched well with mFISH labeling for those cells with that labeling. Specifically, cells with positive PVALB labeling had high PVALB prediction probabilities, whereas cells with positive SST or VIP labeling has low PVALB prediction probabilities (**Figure 6b**). Examples of the intrinsic properties of a cell called as PVALB-positive by the classifier (with confirmed PVALB labeling), and a cell called as non-PVALB (and labeled positive for SST) are shown in **Figure 6c**.

The features with highest weighting in the classifier were AP height, depolarizing sag, AP upstroke adaptation ratio, and membrane time constant (**Supplementary Table 2**). As shown in **Figure 6d**, these features were correlated with the classifier prediction of PVALB vs. non-PVALB identity. Notably, we observed that the separation between PVALB-positive and other interneuron types is not as robust in human recordings compared to mouse, and that appears to be reflected in the somewhat continuous relationship between electrophysiological features and predicted interneurons subclass identity. With this classification of postsynaptic interneurons measured in MPC recordings, we looked at the relationship between synaptic features and their PVALB probabilities (**Figure 6e**). As expected, cells with a high likelihood of being PVALB-positive tended to show synaptic depression as shown by the correlation between paired pulse ratios and PVALB probability using 50 Hz pulse trains. In addition, there was a trend for cells with higher likelihood of being PVALB-positive to have higher EPSP amplitudes, although this was not significant (p = .054). Similar trends were seen with 20 Hz stimulation protocols, although with smaller correlations to PVALB probability that were not significant (**Figure 6 – Figure supplement 1**).

Taken together, this combination of *post-hoc* marker labeling and computational classifier predictions indicate that we can identify postsynaptic cell identity for PVALB versus other, non-PVALB interneuron types in MPC recordings. With these postsynaptic cells identified, this allows a conclusion that synaptic properties between presynaptic human pyramidal neurons and postsynaptic interneurons are target-dependent based on the interneuron subclass identity, with PVALB neurons more likely to show synaptic depression and non-PVALB neurons more likely to show synaptic facilitation. We also observed more facilitating synapses in slice cultures than acute slices. While this could be due to a slice culture artifact that affects dynamics of short-term plasticity, it is also plausible this is the consequence of differential interneuron subclass sampling in the two conditions. Specifically, the percentage of neurons predicted to be non-PVALB neurons in acute slice recordings (10%, 1 out of 10 recorded postsynaptic neurons) was much lower than in slice culture (∼41%, 18 out of 43 recorded postsynaptic neurons). Thus, it is likely that there is a selection bias for PVALB neurons when patching in unlabeled acute slices, and the AAV-based strategy with a pan-GABAergic enhancer allows a more unbiased sampling of interneuron subclasses whose properties are presumably preserved in culture.

## DISCUSSION

### Target cell-dependent excitatory to inhibitory neuron synaptic properties in human cortex

Rodent studies have established that properties of short-term synaptic dynamics excitatory and inhibitory neuron connections in the cortex and other regions are often dependent on postsynaptic neuron identity (Blackman et al., 2013). The current study establishes that this principle is also true in human cortex. Using MPC recordings in human neurosurgically resected cortical tissues, we find that individual pyramidal neurons show heterogeneous synaptic properties to multiple postsynaptic GABAergic neurons, and that those properties are defined by postsynaptic neuron identity. As in mouse, PVALB-positive fast-spiking interneurons tended to show synaptic depression, whereas SST-positive (or more generally, PVALB-negative) interneurons tended to show synaptic facilitation. Given the conserved intrinsic properties of human and rodent PVALB-positive neurons (i.e., fast spiking) and their target-specific depressing synaptic dynamics, PVALB-positive “basket” like cells in human cortex are very likely to have similar functional roles in cortical circuits. These roles likely include mediation of excitation-inhibition balance, gain control, and generation/synchronization of fast oscillation (e.g., “gamma” frequency range, 20-80 Hz) by communicating with reciprocally connected neighboring excitatory neurons (Isaacson and Scanziani, 2011). Non-PVALB interneurons, including SST-stained neurons by *post-hoc* HCR, instead showed rather small initial EPSP responses and tended to have short-term synaptic facilitation. These properties are comparable to previous studies in rodent SST-positive “Martinotti “cells, which are known to target pyramidal neuron apical dendritic tufts and mediate lateral disynaptic inhibition (Silberberg and Markram, 2007; Berger et al., 2009). Putative “Martinotti” cells in human cortex also contribute to lateral disynaptic inhibition between two neighboring pyramidal neurons *via* receiving delayed facilitating synapses (Obermayer et al., 2018). Therefore, both the target cell-dependent principles and the specific properties of synaptic plasticity in PVALB versus other interneuron types appear to be strongly conserved, suggesting similar roles in functional cortical circuitry across species.

Transcriptomic analysis of interneuron subclasses also strongly suggested that synaptic properties would vary by interneuron subclass. Not only are many synaptic genes differentially expressed between subclasses, but subclasses can be identified by *de novo* clustering using only synapse-associated genes. This result suggests that such genes are particularly important for cell identity and function (Paul et al., 2017; Huang & Paul, 2019; Smith et al., 2019), as highlighted by several prior transcriptome studies. While currently challenging to interpret at a gene-to-synaptic function level, since so many synapse -associated genes vary by cell subclass, eventually these molecular data will provide a mechanistic substrate for cell type-specific functional properties and allow prediction of both conserved and divergent properties. For example, as mentioned above *Elfn1* has been shown to control short-term facilitation in SST interneurons cells (Sylwestrak & Ghosh, 2012; de Wit & Ghosh, 2016; Stachniak et al., 2019). We find that *ELFN1* is expressed in all interneuron subclasses except PVALB neurons, and this pattern is conserved from mouse to human, suggesting a similar role in non-PVALB expressing interneurons from mouse to human. On the other hand, prior studies in human cortical tissues have shown a variety of differences from mouse, such as in excitatory neuron recovery from synaptic depression (Testa-Silva et al., 2014), higher presynaptic release probabilities (Testa-Silva et al., 2014), more docked vesicles (Molnar et al., 2016), and polysynaptic network activities (Molnar et al., 2008; Szegedi et al., 2017; Campagnola et al., 2022), as well as diverse forms of synaptic plasticity among specific interneuron types (Verhoog et al., 2013; Szegedi et al., 2016; Mansvelder et al., 2019). Many other synapse-associated genes show differential expression across interneuron subclasses but with divergent expression across species, perhaps underlying these described functional synaptic species differences and predicting that many other details may vary across species.

### Strategies for cell type-specific analysis of synaptic connectivity in human tissues

Directly analyzing synaptic properties of specific connected cell types by MPC experiments in human or other non-genetically tractable model organisms presents a number of challenges. The first challenge is simply access to healthy human tissues for slice physiology experiments. We and others have demonstrated that tissues from human neurosurgical resections are highly robust and can be used both for acute recordings and slice culture experiments over several weeks to months (Eugene et al, 2014; Schwarz et al., 2017; Ting et al, 2018; Berg et al., 2021). Another major challenge is efficiently targeting specific cell types. Typically, human tissue slice physiology is performed in unlabeled tissues with cell type targeting based only on their soma and proximal dendritic shapes under the microscope (Molnar et al., 2008; Molnar et al., 2016). We have taken advantage of the longevity of human slices in slice culture to transduce neurons with enhancer-AAVs to allow viral transgenesis and genetic manipulation of cells in brain slices (Andersson et al, 2016; Le Duigou et al, 2018; Ting et al., 2018; Mich et al., 2021; Schwarz et al, 2019), in the current study to target GABAergic interneurons. Finally, another major challenge is the *post-hoc* identification of recorded neurons in MPC experiments. We demonstrate two effective strategies for cell type identification. The first is a low throughput but high confidence FISH staining of recorded neurons with markers of interneuron subclasses. The second is a quantitative classifier to differentiate interneuron subclass identities based solely on electrophysiology data, using a high confidence Patch-seq dataset that links physiology with transcriptomic identity to build the classifier. Together this array of solutions allowed the conclusions to be drawn about target cell-dependent synaptic properties at the GABAergic subclass level, and these approaches should be possible to apply at a much finer level of cell type resolution in the future.

A key strategy demonstrated here is to use mFISH with multiple rounds of staining on cleared thick *in vitro* human slice preparations, preserving tissue integrity and cell morphology, thereby allowing molecular identification of synaptically connected neurons using robust marker genes for neuron subclasses. The use of mFISH provides advantages over traditional immunohistochemical staining, as unambiguous identification of interneuron subclass identity (e.g., PVALB, SST, VIP, LAMP5) has not been reliable with *post-hoc* immunohistochemical staining in both non-human primate and human tissues. For example, PVALB antibodies work well (Szegedi et al., 2017), but SST and VIP antibodies do not work reliably in human cortical slices in our hands (data not shown; but see Lukacs et al., 2022). Here, the GABAergic interneuron subclasses PVALB, SST, and VIP were readily resolved using HCR and RNA transcript probes for *PVALB*, *SST*, *VIP* and *LAMP5*. However, several challenges were identified for future improvement. Although mRNA labeling was robust for abundant genes, less abundant genes were more difficult to detect. Autofluorescence from lipofuscin, a common feature of human brain tissue, can complicate analysis and obscure mRNA signal. Improvement of lipofuscin mitigation techniques will facilitate future analysis. In some cases, we did not readily detect expected mRNA transcripts for cells with certain types of electrophysiological features (such as fast-spiking inhibitory neurons that would be expected to express PVALB). This could be the true state of the cell, or due to loss of mRNA through the patch pipette or leakage from the cell after pipette withdrawal in addition to HCR based gene detection and amplification procedures in thick human surgical tissues. Finally, greater cell type resolution will be gained using more highly multiplexed mFISH techniques (Chen et al., 2015; Eng et al., 2019; Wang et al., 2021). Despite these opportunities for further refinement, the approach utilized here enables multiple modality functional interrogation of human brain cell types and is a valuable step toward deciphering the correspondence between these multiple data modalities.

The development of AAV vectors for rapid infection and cell type-specific transgene expression, provides new avenues for targeted analysis in the human brain as well as in non-genetically tractable organisms. This method provides a means to study neuronal and circuit properties in human neocortex and link them to emerging molecular classifications of cell types (Tasic et al., 2018; Hodge et al., 2019; Bakken et al., 2021). Efforts at identification of cell type-selective genomic enhancers are quickly generating new enhancer-AAV tools to target a wide variety of cortical cell types (Graybuck et al., 2021; Mich et al., 2021). There are clear advantages to use novel viral tools to prospectively label specific cell subclasses in a slice culture platform and investigate their intrinsic membrane properties and synaptic connectivity during MPC recordings in a targeted manner. However, there are two things that need to be carefully considered: one is potential cultural artifact (Ting et al., 2018; Suriano et al., 2021) and the other is potential modification of synaptic properties in virally labeled neurons. Therefore, data obtained from virally labeled slice cultures ultimately need to be compared to data extracted from acute slice (Ting et al., 2018; Schwarz et al., 2019). Nonetheless, genetic labeling of rare cell types will be very helpful to investigate intrinsic properties and synaptic connectivity, and offers potential for cell type-specific functional manipulation in mature human brain tissues.

## METHODS

### Transcriptomic analysis

Previously described single nucleus transcriptomic datasets from human MTG (Hodge et al., 2019) and mouse VISp (Tasic et al., 2018) were analyzed to define differentially expressed genes between PVALB and SST neuron types. Expression matrices were reduced to 14,870 orthologous genes conserved between human and mouse. A differential expression analysis between PVALB and SST subtypes was performed on log2 normalized data using the ‘FindMarkers’ function in Seurat v4.0.4 (Hao et al., 2021) with the Wilcoxon rank sum test. Genes were defined as differentially expressed if their log2 fold change was greater than 0.5 and their adjusted p-value was less than 0.01. Genes that were differentially expressed for PVALB or SST types in both species were used for heatmaps and gene ontology analysis (**Supplementary Table 1**). For gene ontology analysis, the gene universe was defined by orthologous genes that had greater than 0 expression in PVALB or SST nuclei. The ‘enrichGO’ function from R package clusterProfiler (Wu et al., 2021; Yu et al., 2012) was used to compare conserved PVALB and SST DEG lists to the gene universe background with Benjamini-Hochberg correction and pvalueCutoff set to 0.01 and qvalueCutoff set to 0.05. Enriched terms were ranked by adjusted p-value and the top 10 terms for cellular compartment were shown.

**Supplementary Table 1.**
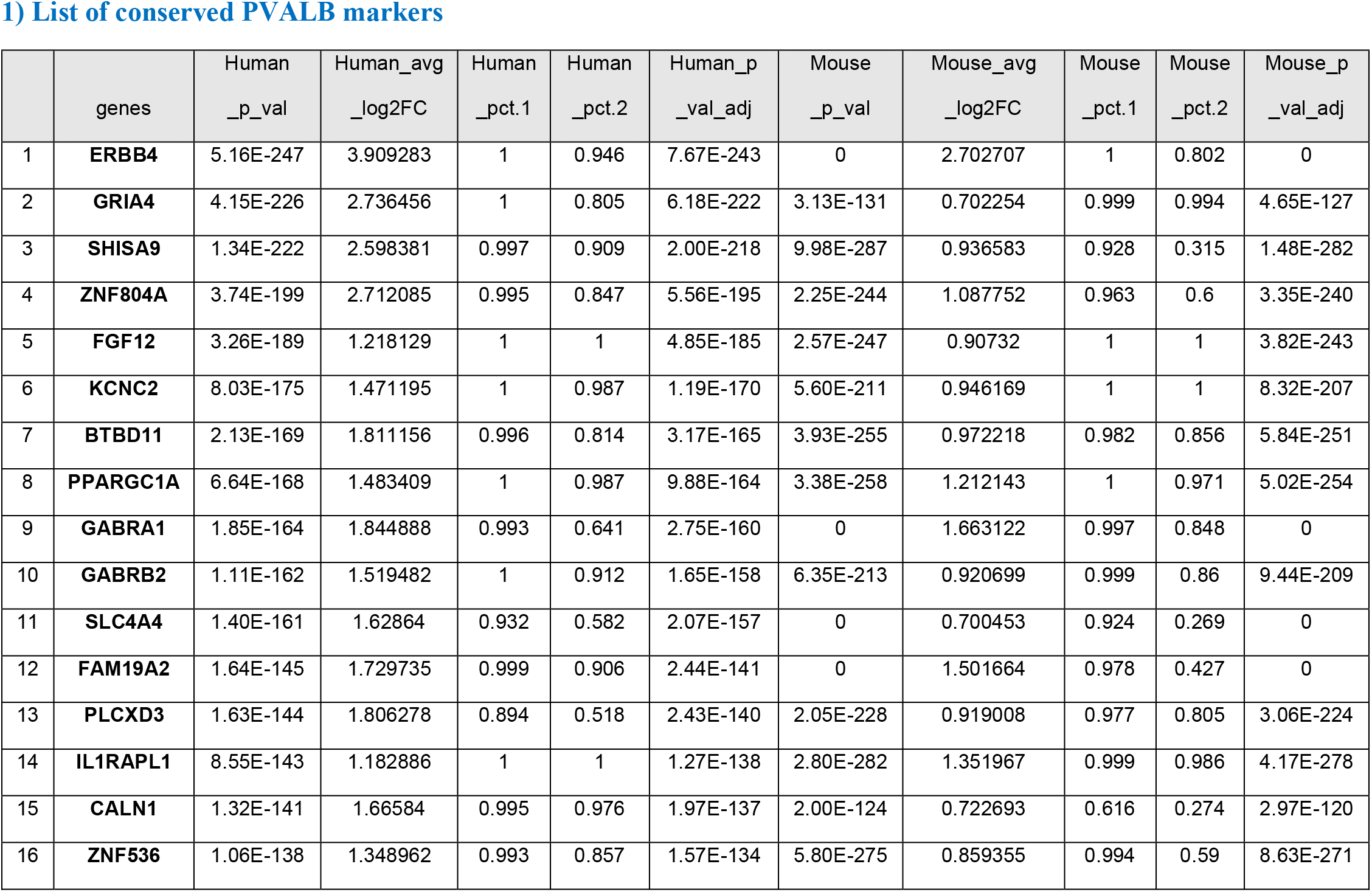
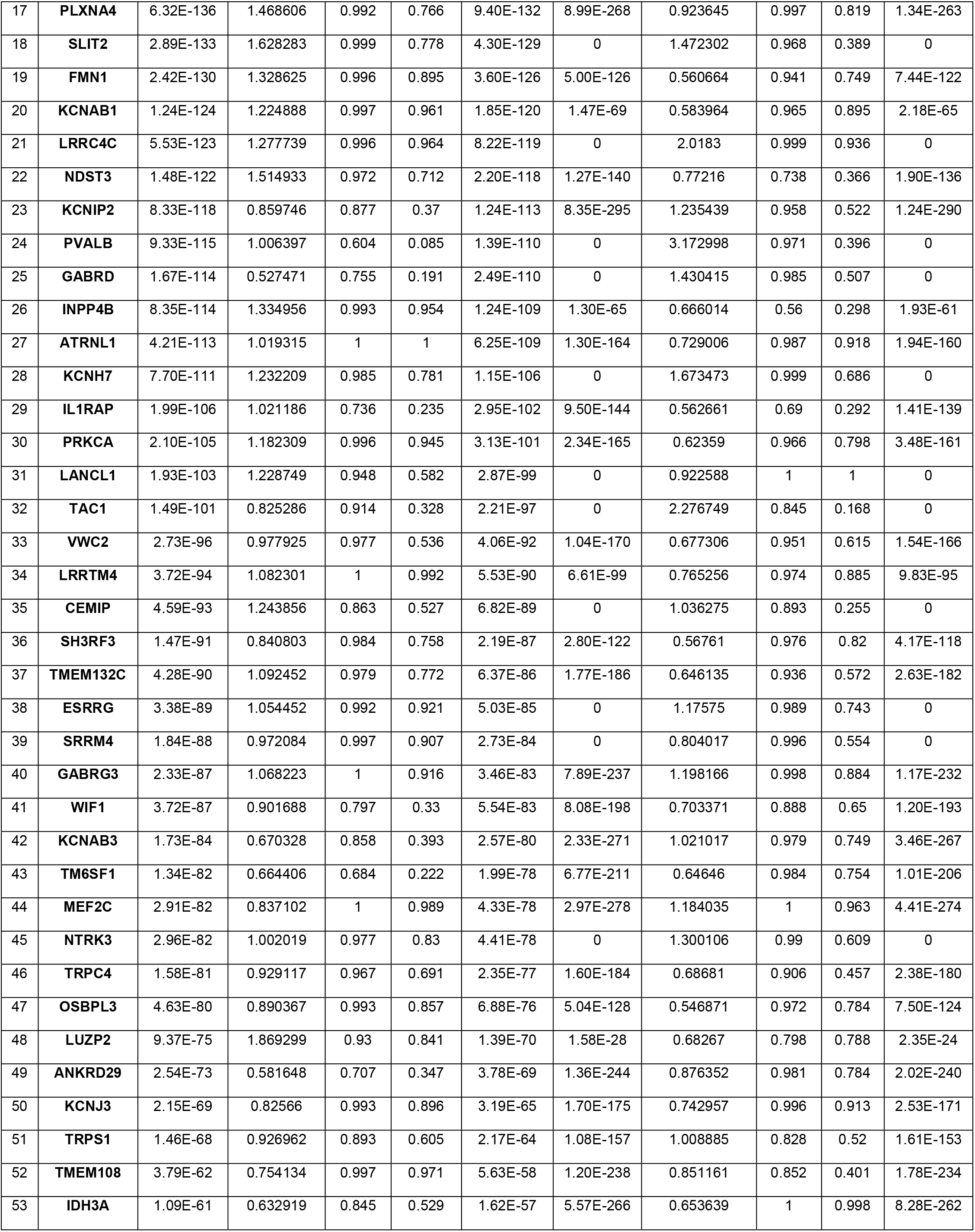
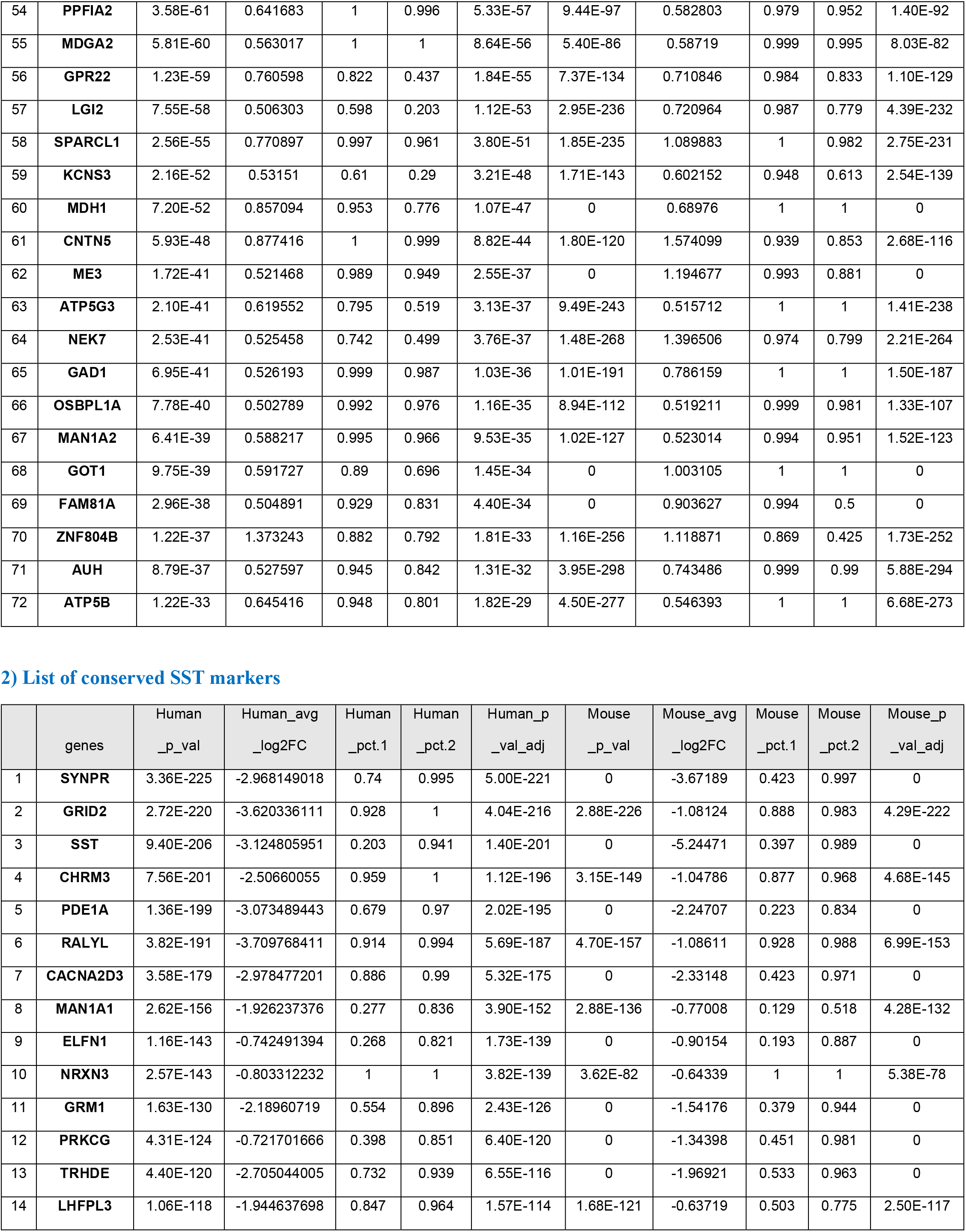
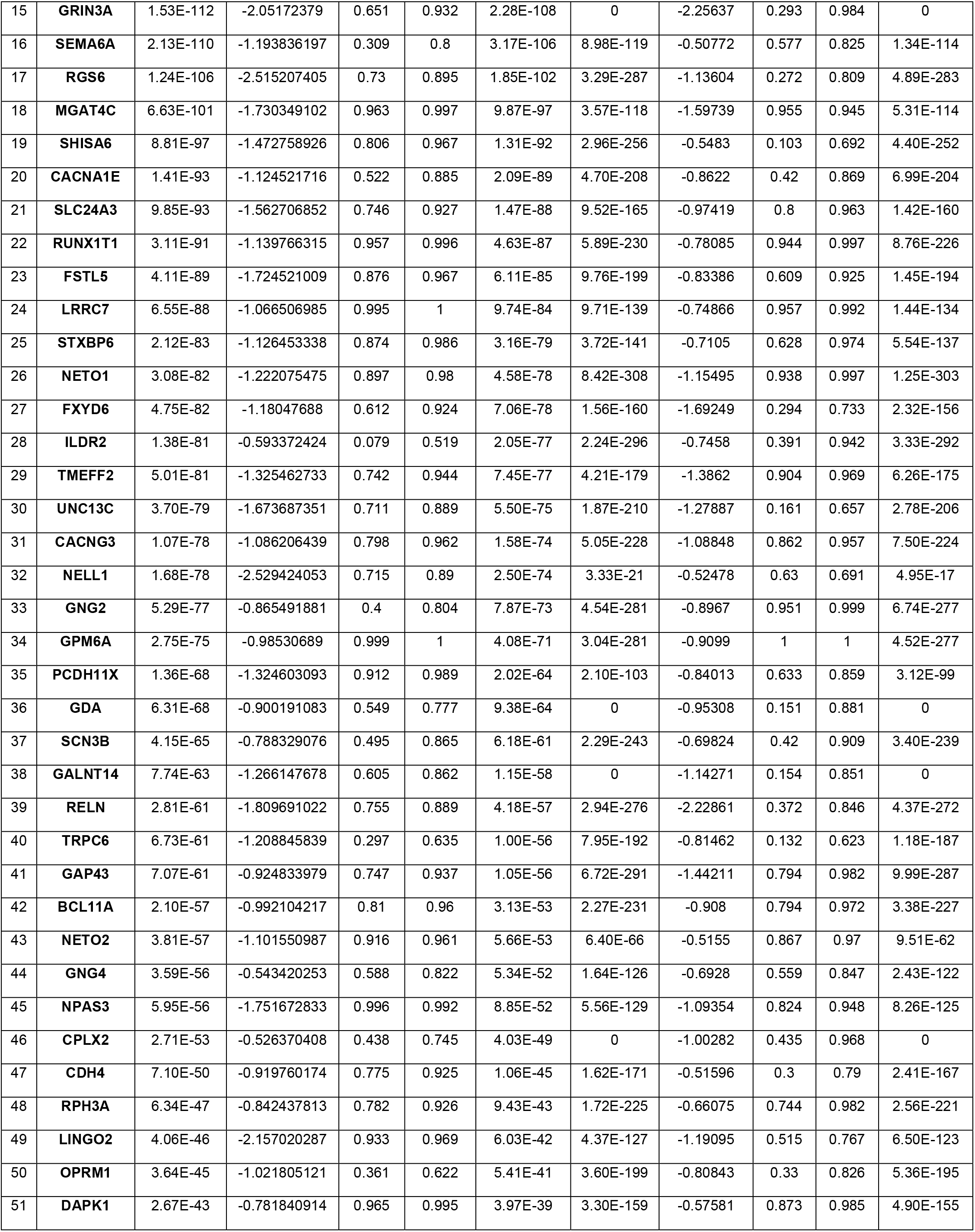
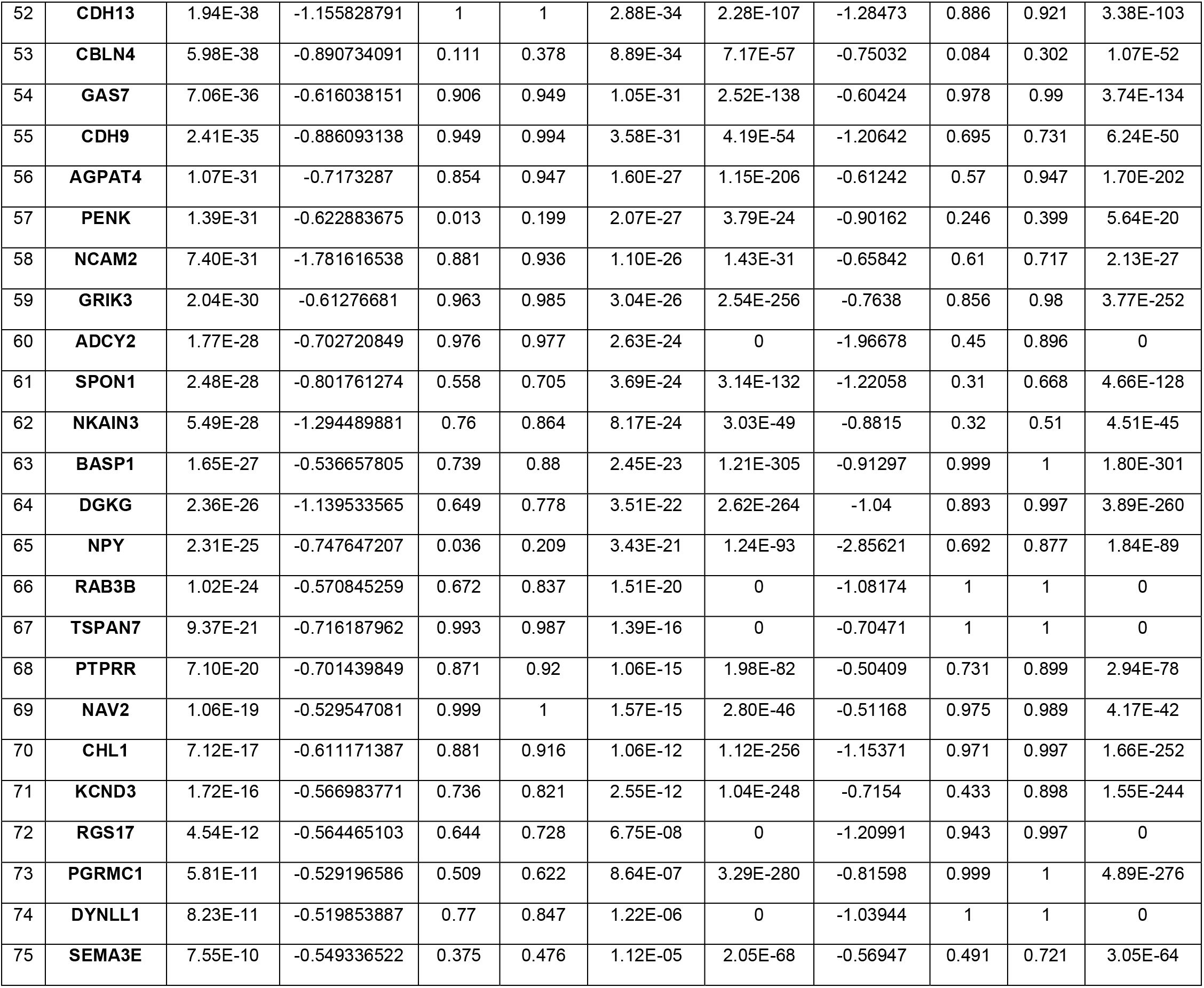
PVALB versus SST subclass differential gene expression analysis. p_val and p_val_adj indicate significance and adjusted significance of the differential expression test (Wilcoxon sum rank test). avg_log2FC means the average log2 fold change in expression between the two cell populations (pvalb and sst). pct.1 is the proportion of target nuclei a gene is expressed in, and pct.2 the proportion of the background population the gene is expressed in.

### Acute slice preparation

Human cortical tissues were collected from adult patients undergoing neurosurgical procedures to treat symptoms associated with either epilepsy or brain tumor. Surgical specimens were obtained from local hospitals (Harborview Medical Center, Swedish Medical Center, and University of Washington Medical Center) in collaboration with local neurosurgeons. Surgically resected neocortical tissue was distal to the pathological core (i.e., tumor tissue or mesial temporal structures). Detailed histological assessment and using a curated panel of cellular marker antibodies indicated a lack of overt pathology in surgically resected cortical slices (Berg et al., 2021). In this study, we included data from 31 surgical cases, 15 of which were epilepsy cases and the remaining 16 were tumor cases (**Figure 2 – Figure supplement 1**). All specimens derived from neocortex with the majority of cases derived from the temporal cortex (*n* = 21) while a minority were obtained from the frontal cortex (*n* = 9) or anterior cingulate cortex (*n* = 1).

Surgical specimens were immediately transported (15-35 min) from the operating room to the laboratory in chilled (0-4°C) artificial cerebral spinal fluid (aCSF) slicing solution containing (in mM): 92 N-Methyl-D-glucamine (NMDG), 2.5 KCl, 1.25 NaH_2_PO_4_, 30 NaHCO_3_, 20 4-(2-hydroxyethyl)-1-piperazineethanesulfonic acid (HEPES), 25 D-glucose, 2 thiourea, 5 Na-L-ascorbate, 3 Na-pyruvate, 0.5 CaCl_2_, and 10 MgSO_4_ (Ting et al., 2014). The NMDG aCSF was continuously bubbled with carbogen (95% O_2_ and 5% CO_2_). Osmolality was measured and adjusted to 300-315 mOsmoles/kg range (305-315 mOsmoles/kg range when using a freezing point osmometer, and 300-310 mOsmoles/kg range if using vapor pressure osmometer), and the pH was measured and adjusted to 7.3-7.4. 350 µm thick human cortical slices were prepared using a Compressome VF-300 (Precisionary Instruments) or VT1200S (Leica Biosystems). After being cut, slices were transferred to oxygenated NMDG aCSF maintained at 34°C for 10 min. Slices were kept at room temperature in oxygenated holding aCSF solution containing (in mM): 92 NaCl, 30 NaHCO_3_, 25 D-Glucose, 20 HEPES, 5 Na-L-Ascorbate, 3 Na Pyruvate, 2.5 KCl, 2 CaCl_2_, 2 MgSO_4_, 2 Thiourea, 1.2 NaH_2_PO_4_ prior to recording (Seeman et al., 2018; Berg et al., 2021; Lee et al., 2021; Campagnola, Seeman et al., 2022).

### Slice culture preparation

Following brain slice preparation and NMDG recovery steps as outlined above, a subset of brain slices were transferred to a 6-well plate for culture and viral transduction. Human cortical brain slices were placed on membrane inserts (Millipore #PICMORG), and the wells were filled with 1 mL of culture medium consisting of 8.4 g/L MEM Eagle medium, 20% heat-inactivated horse serum, 30 mM HEPES, 13 mM D-glucose, 15 mM NaHCO_3_, 1 mM ascorbic acid, 2 mM MgSO_4_, 1 mM CaCl_2_, 0.5 mM GlutaMAX-I, and 1 mg/L insulin (Ting et al 2018). The slice culture medium was carefully adjusted to pH 7.2-7.3, osmolality of 300-310 mOsmoles/Kg by addition of pure H_2_O, sterile-filtered and stored at 4°C for up to two weeks. Culture plates were placed in a humidified 5% CO_2_ incubator at 35°C. 1-3 hours after brain slices were plated on cell culture inserts, brain slices were infected by direct application of concentrated AAV viral particles over the slice surface (Ting et al 2018). The slice culture medium was replaced every 2-3 days until initiating synaptic physiology experiments. The time window to perform slice culture experiments ranged from 2.5 to 9 DIV, and a total of 36 cultured human neocortical slices were used in this study for the identification of gene expression with mFISH/HCR after MPC recordings.

### Viral vector production

Recombinant AAV vectors were produced by triple-transfection of ITR-containing enhancer plasmids along with AAV helper and rep/cap plasmids using the AAV293 cell line, followed by harvest, purification and concentration of the viral particles. The AAV293 packaging cell line and plasmid supplying the helper function are available from a commercial source (Cell Biolabs). The PHP.eB capsid variant was generated by Dr. Viviana Gradinaru at the California Institute of Technology (Chan et al., 2017) and the DNA plasmid for AAV packaging is available from Addgene (plasmid#103005). Quality control of the packaged AAV was determined by qPCR to determine viral titer (viral genomes/mL), and by Sanger sequencing of the AAV genome to confirm the identity of the viral vector that was packaged.

### CN1390 vector design and construction

Human neocortical interneurons were targeted in cultured slices by transducing slices with an optimized forebrain GABAergic viral vector CN1390, also known as pAAV-DLX2.0-SYFP2. The DLX 2.0 sequence includes a 3x concatemer of the core region of a previously well-characterized DLX I56i forebrain GABAergic neuron enhancer (Dimidischstein et al 2016; Zerucha et al, 2000). The 131 bp core sequence of the hI56i enhancer was inferred from enhancer bashing experiments detailed in Zerucha et al, 2000. The 393 bp 3x core enhancer concatemer sequence was custom gene synthesized and subcloned into pAAV-minBetaGlobin-SYFP2-WPRE3-BGHpA upstream of the minimal promoter to make pAAV-DLX2.0-SYFP2, vector ID# CN1390 in our catalog. This vector will be deposited to Addgene for distribution to the academic community upon publication.

### Electrophysiology

Experiments were conducted on an upright microscope with an oblique condenser (WI-OBCD, Olympus) equipped with infrared (850 nm) illumination, 490 nm, 565 nm and ultraviolet laser (395 nm) lines (Thorlab). 4x and 40x objectives (Olympus) were used to visualize the sample and a digital CMOS camera (Flash 4.0 V2, Hamamatsu) to take images. The rig configuration included eight electrodes disposed around the recording chamber, each surrounded by an headstage shield in order to prevent electrical crosstalk artifacts. Each patch electrode was positioned by x-y stage and micromanipulator (PatchStar, Scientifica) with guidance of acq4 open python platform software (acq4.org; Campagnola et al., 2014). Bright-field and fluorescent images were also captured and analyzed with acq4. Signals were amplified using Multiclamp 700B amplifiers (Molecular Devices) and digitized at 50-200 kHz using ITC-1600 digitizers (Heka). Pipette pressure was controlled using electro-pneumatic pressure control valves (Proportion-Air; PA2193). The recording software, Igor Pro7 or 8 (WaveMetrics), contained with a custom software Multi-channel Igor Electrophysiology Suite (MIES; https://github.com/AllenInstitute/MIES), used to apply the bias current, inject the appropriate amount of current to patched cells, data acquisition and pressure regulation.

Slices were transferred to the recording chamber and perfused with carbogenated aCSF (2mL/min), constant temperature (31-32 °C), pH 7.2-7.3 and oxygen saturation in the recording chamber (40-50%). Perfusing aCSF contained (in mM): 1.3 CaCl_2_, 12.5 D-Glucose, 1 or 2 MgSO_4_, 1.25 NaH_2_PO_4_, 3 KCl, 18 NaHCO_3_, 126 NaCl, 0.16 Na-L-Ascorbate. Patch pipettes were pulled from thick-wall filamented borosilicate glass (Sutter Instruments) using a DMZ Zeitz-Puller (Zeitz) to a tip resistance of 3-8 MΩ, and filled with intracellular solution containing (in mM) either 0.3 ethylene glycol-bis(b-aminoethyl ether)-N,N,N’,N’-tetraacetic acid (EGTA) or no EGTA in addition to: 130 K-gluconate, 10 HEPES, 3 KCl, 0.23 Na_2_GTP, 6.35 Na_2_Phosphocreatine, 3.4 Mg-ATP, 13.4 Biocytin, and fluorescent dye with 50 µM Alexa-488 or cascade blue. Solution osmolarity ranged from 280 to 295 mOsmoles/kg titrated with sucrose, pH between 7.2 and 7.3 titrated with KOH. The liquid junction potentials were not corrected. For slice culture experiments, GABAergic neurons labeled with AAV-DLX2.0-SYFP2 were targeted during patch pipettes were approaching. With cascade blue loaded in the patch pipette, overlaid signals in the same cells with both SYFP2 and cascade blue were confirmed by manual inspection of image stacks with blue and green LED light excitation.

Cell cluster of eight neurons at each trial was selected and attempt for multiple whole-cell patch-clamp (MPC) recordings, targeted in mainly supraganular layer (L2 and L3), 50-80 µm depth from slice surface and smooth somatic appearance. Pairwise recordings were performed for local synaptic connectivity assay with both voltage and current-clamp mode. In voltage-clamp mode, membrane voltages of all patched cells were hold at either −70 or −55 mV and brief depolarization to 0 mV for 3 ms at 20 Hz sequentially to reliably identify both excitatory and inhibitory connections. In current-clamp mode, initially all cell membrane potentials were maintained at −70 ± 2 mV with automated bias current injection when we generated presynaptic unitary action potential by brief current injections (1.5-3 ms) to detect EPSP responses in postsynaptic cells. For inhibitory connection, cell membrane potentials were maintained at −55 ± 2 mV to detect IPSP responses in postsynaptic cells.

For the short-term plasticity, there are 12 action potentials at multiple frequencies (10, 20, 50, and 100 Hz) to induce sequential postsynaptic responses in connected pairs. Presynaptic stimulus amplitudes were adjusted to generate unitary action potential in each pulse. In order to measure recovery time course after induction protocol (i.e., initial 8 pulses), inter-spike interval between 8^th^ and 9^th^ pulses at 50 Hz stimulation was varied sequentially at 62.5, 125, 250, 500, 1000, 2000, and 4000 ms. For other frequency stimulation (10, 20, and 100 Hz), we used fixed 250 ms inter-spike interval between 8^th^ and 9^th^ pulses. Stimuli were interleaved between cells such that only one cell was spiking at a time, and no two cells were ever evoked to spike within 150 ms of each other (Seeman et al., 2018; Campagnola, Seeman et al., 2022). At each sequential 12 pulses stimulation for all patched neurons were repeated with 15 s inter-sweep interval. After running connectivity protocol, step current injections in each cell were applied to extract intrinsic membrane properties such as spike shape and frequency-current relationship.

### Human cortical interneuron patch-seq recordings in virally labeled slice cultures

Similar experimental procedures were applied as described in previous studies (Berg et al., 2021; Lee et al., 2021).

Slices were bathed in warm (32-34 °C) recording aCSF containing the following (in mM): 126 NaCl, 2.5 KCl, 1.25 NaH_2_PO_4_, 26 NaHCO_3_, 12.5 D-glucose, 2 CaCl_2_.4H_2_O) and 2 MgSO_4_.7H_2_O (pH 7.3), continuously bubbled with 95% O_2_ and 5% CO_2_. The bath solution contained blockers of fast glutamatergic (1 mM kynurenic acid) and GABAergic synaptic transmission (0.1 mM picrotoxin).

Recording pipettes were filled with ∼1.75 μL of RNAse Inhibitor containing internal solution: 110 mM K-Gluconate, 4 mM KCl, 10 mM HEPES, 1 mM adenosine 5’-triphosphate magnesium salt, 0.3 mM guanosine 5’- triphosphate sodium salt hydrate, 10 mM sodium phosphocreatine, 0.2 mM ethylene glycol-bis (2-aminoehtylether)-N,N,N’,N’-tetraacetic acid, 20 μg/mL glycogen, 0.5 U/μL RNase Inhibitor, 0.5 % biocytin, and either 50 μM Cascade Blue dye (excited at 490 nm), or 50 μM Alexa-488 (excited at 565 nm).

After examination of intrinsic membrane properties of virally labeled interneurons in conventional patch-clamp recordings, a small amount of negative pressure was applied (∼-30 mbar) to extract the nucleus to the tip of the pipette. After extraction of the nucleus, the pipette containing internal solution, cytosol, and nucleus was removed from the pipette holder and contents were transferred into a PCR tube containing lysis buffer. CDNA amplification, library construction, and subsequent RNA-sequencing procedures are described in Berg et al., 2021 and Lee et al., 2021. Patch-seq data was mapped to the reference taxonomy from human MTG dissociated cells (Hodge et al., 2019; Gouwens et al., 2020).

### Classification of intrinsic membrane properties in postsynaptic interneurons against a reference dataset obtained from single-cell patch-seq data

Intrinsic characterization of individual cells from both acute and slice cultures was carried out as described in (Campagnola, Seeman et al., 2022). Features were primarily calculated from sweeps with long square pulse current injection: subthreshold properties such as input resistance, sag, and rheobase; spike train properties such as f-I slope and adaptation index; and single spike properties such as upstroke-downstroke ratio, after-hyperpolarization, and width. For spike upstroke, downstroke, width, threshold, and inter-spike interval (ISI), ‘adaptation ratio’ features were calculated as a ratio of the spike features between the first and third spike. A subset of cells also had subthreshold frequency response characterized by a logarithmic chirp stimulus (sine wave with exponentially increasing frequency), for which the impedance profile was calculated and characterized by features including the peak frequency and peak ratio. Feature extraction was implemented using the IPFX python package (https://github.com/AllenInstitute/ipfx); custom code used for chirps and some high-level features will be released in a future version of IPFX.

Prediction of cell subclass from intrinsic properties was accomplished using a classifier trained against a reference dataset of cells with intrinsic properties and known subclasses. Reference cells were targeted in human slice culture with the same enhancer, AAV-DLX2.0-SYFP2 as the primary dataset, gene expression characterized using the patch-seq protocol (Berg et al., 2021; Lee et al., 2021), and transcriptomic subclasses assigned by mapping to a reference transcriptomic taxonomy from (Hodge et al., 2019) following the method from (Gouwens et al., 2019). However, differences in recording conditions between synaptic physiology and patch-seq protocols (primarily the presence/absence of synaptic blockers) cause shifts in intrinsic properties that preclude the use of all features in this reference dataset. We therefore excluded features for which the protocol accounted for over 5% of variance in an ANOVA of the combined dataset, leaving 24 features but excluding some common discriminating features such as spike width. Using scikit-learn, we trained a classifier pipeline that first normalizes features based on a robust variance (RobustScaler), imputes missing values based on nearest neighbors (KNNImputer), then classifies with linear discriminant analysis. This pipeline achieved 77% accuracy for predicting PVALB/non-PVALB subclasses in the reference dataset using cross-validation. Errors came primarily from a subset of SST cells with intrinsic properties overlapping the PVALB cells. The PVALB prediction probabilities of the classifier were then calibrated on the full reference dataset (CalibratedClassifierCV) before applying to the synaptic physiology dataset to generate both PV probabilities. Cells with a low confidence (PVALB probability between 0.4 and 0.6) were categorized as uncertain, with higher probabilities labeled PVALB and lower labeled non-PVALB. The separation of subclasses and overlap between patch-seq reference and MPC cells in intrinsic feature space was visualized using Uniform Manifold Approximation and Projection, UMAP (Becht et al., 2018).

**Supplementary Table 2.**
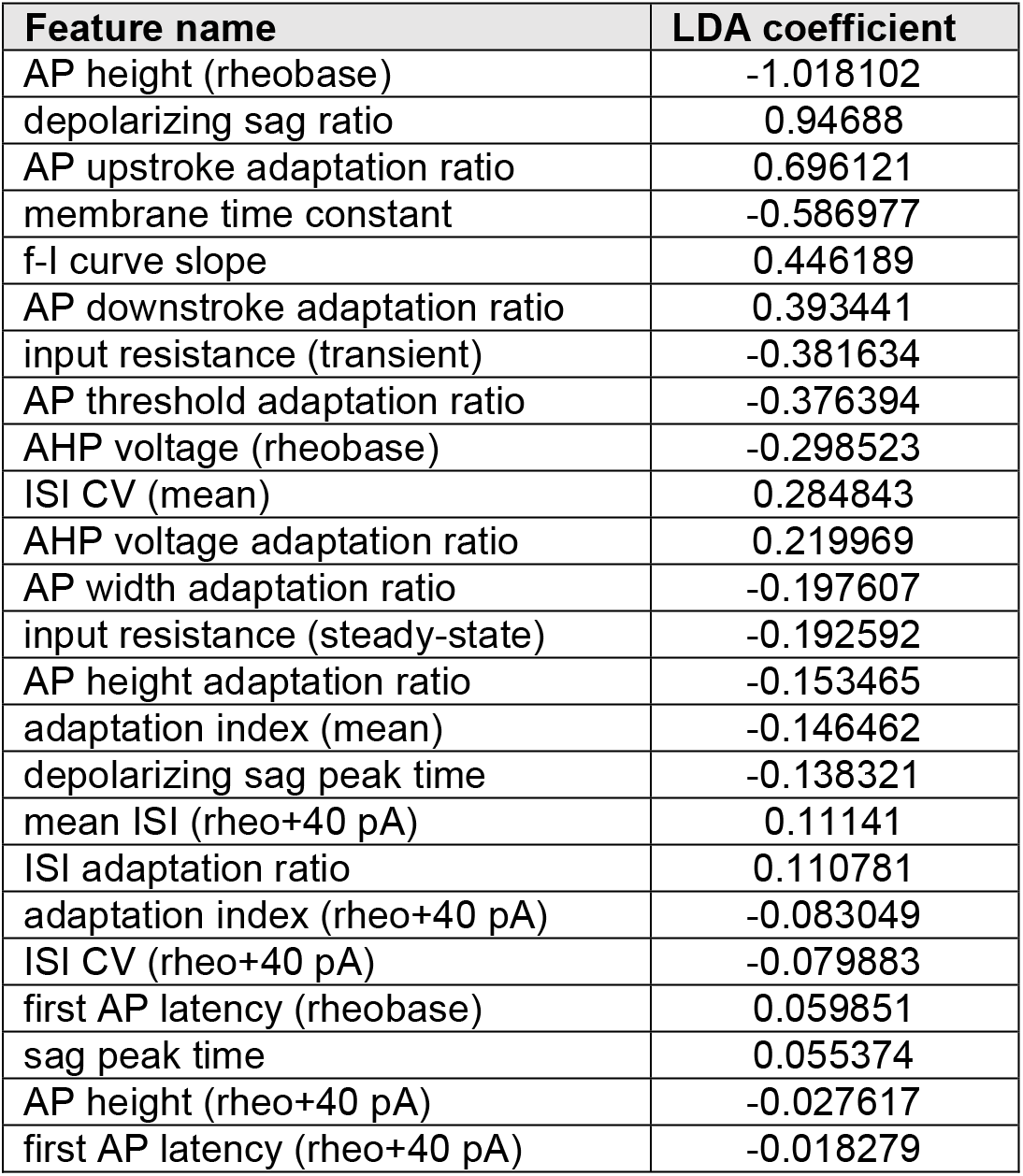
List of intrinsic membrane properties classifier features and linear discriminant analysis (LDA) coefficients.

### Data analysis

Synaptic connectivity and dynamics, intrinsic membrane properties were analyzed with custom-written MATLAB (MathWorks) and Igor (Wavemetrics) software. Somatic position of individual neurons in a cluster from electrophysiological recording was imaged with fluorescent dyes (Alexa488 or cascade blue) with upright microscope and saved in ACQ4. Consequently, recorded neurons were identified with biocytin staining image and matched with mFISH/HCR signals taken by inverted confocal microscope.

To determine whether presynaptic spike generation is intact by a brief somatic current injection, all recorded presynaptic traces were manually checked and quality controlled based on the spike shape. When presynaptic spike shape is intact, postsynaptic response failures were included to average EPSP responses with multiple stimulations. EPSP onset delay was calculated from the peak of presynaptic spikes in current clamp mode to the onset of EPSP response. EPSP onset delay, PSP rise time, PSP decay tau were calculated with some modification of codes from Postsynaptic Potential Detector shared in public (MATLAB Central File Exchange; https://www.mathworks.com/matlabcentral/fileexchange/19380-postsynaptic-potential-detector, 2020).

### Thick tissue mFISH sample preparation

Slices were fixed in 4% PFA for 2 hours at room temperature (RT), washed three times in PBS for 10 min each, then transferred to 70% EtOH at 4°C for a minimum of 12 hours, and up to 30 days. Slices were then incubated in 8% SDS in PBS at RT for two hours with agitation. The solution was exchanged with 2X sodium chloride sodium citrate (SSC) three times, slices were washed for one hour at RT, followed by two additional (1 hour each) washes with fresh 2X SSC.

### *In situ* HCR for thick tissue

We performed HCR v3.0 using reagents and a modified protocol from Molecular Technologies and Molecular Instruments (Choi et al., 2014). Slices were incubated in pre-warmed 30% probe hybridization buffer (30% formamide, 5X sodium chloride sodium citrate (SSC), 9 mM citric acid pH 6.0, 0.1% Tween 20, 50 µg/mL heparin, 1X Denhardt’s solution, 10% dextran sulfate) at 37°C for 5 min, then incubated overnight at 37°C in hybridization buffer with the first three pairs of probes added at a concentration of 4 nM. The hybridization solution was exchanged 3 times with 30% probe wash buffer (30% formamide, 5X SSC, 9 mM citric acid pH 6.0, 0.1% Tween 20, 50 µg/mL heparin) and slices were washed for one hour at 37°C. Probe wash buffer was briefly exchanged with 2X SSC, then amplification buffer (5X SSC, 0.1% Tween 20, 10% dextran sulfate) for 5 min. Even and odd hairpins for each of the three genes were pooled and snap-cooled by heating to 95°C for 90 seconds then cooling to RT for 30 min. The hairpins were then added to amplification buffer at a final concentration of 60 nM, and slices were incubated in amplification solution for 4 hours at RT. This was followed by a brief wash with 2X SSC and a one hour, room temperature incubation in 2X SSC containing 8 µg/µl Brilliant Violet 421TM Streptavidin (BioLegend, Cat. No. 405225) and 0.05% Tween 20. Slices were washed three times for 10 min in 2X SSC. For each round of imaging, an aliquot of 67% 2,2’-Thiodiethanol (TDE) solution was prepared for use as a clearing and immersion fluid. ≥99% TDE (Sigma-Aldrich) was mixed with DI water to create a 67% TDE solution with a refractive index of 1.46, verified by a pocket refractometer (PAL-RI, Atago). Slices were transferred to 67% TDE and allowed to equilibrate for at least 1 hour at room temperature prior to imaging.

### Quantification of thick tissue mFISH signals

Patched cells from acute and cultured tissues were hand segmented volumetrically using QuPath software (Bankhead et al., 2017). Segmentation was performed on either the SYFP2 labeled cell body (slice culture preparation) or HCR signal (acute slice preparation) in transcript positive cells. Additionally, several nearby cells were also segmented in order to characterize typical expression levels in each probed gene and to compare signal level to patched cells. For each imaged channel, a histogram of non-cellular pixels was used to calculate a background threshold, which was taken to be three times the half width at half maximum above median of the distribution of pixel values. A mask of lipofuscin pixels was constructed by first taking all pixels that exceeded this threshold in all HCR channels. This mask was additionally expanded by morphological dilation with a kernel of radius one pixel, iterated two times. For each segmented cell, this mask was applied to each channel and the remaining intensity above background was integrated and normalized to the cell volume, this is taken as a measure of expression in each channel and reported in **Figures 3,10** and **Supplementary Figures 3,4,5**.

### Confocal imaging

Thick tissue images were acquired on an Olympus FV3000 confocal microscope using a 30X silicon oil objective with the zoom set to 1.5x. The image montage stacks were acquired through the depth of the tissue at 1.2 µm steps. For figures, maximum intensity projections though the region of interest were generated are shown. Note that some montages exhibit stitching artifacts. Due to the frequent appearance of lipofuscin in aging human tissues, we showed HCR images as multiple overlapping channels since the lipofuscin granules were revealed as spots that are fluorescent in every channel.

### Stripping and subsequent hybridization rounds

To strip the signal in preparation for subsequent rounds, 67% TDE was exchanged with 2X SSC three times and samples were washed for 1 hour. 2X SSC was replaced with 1X DNase buffer for 5 min and then a 1:50 dilution of DNase I in DNase buffer (DNase I recombinant, RNase-free, Roche, Cat. No. 04716728001), and incubated for 1 hour at 37°C. This solution was replaced with fresh DNase solution before incubating slices overnight at 37°C. Slices were washed with 65% formamide in 2X SSC for one hour at 37°C, then 2X SSC for one hour at RT, before being transferred to 67% TDE for at least one hour. After imaging to confirm the signal was gone, the slices were washed in 2X SSC for one hour to remove TDE before proceeding to subsequent hybridization rounds, which followed the protocol described above, except omitting the incubation in streptavidin solution.

### Morphological reconstruction

Reconstructions of the dendrites and the initial part of the axon (spiny neurons) and/or the full axon (aspiny/sparsely spiny neurons) were generated for a subset of neurons with good-quality electrophysiology and biocytin fills. Reconstructions were generated based on a 3D image stack taken by confocal microscope that was run through a Vaa3D-based image processing and reconstruction pipeline (Peng et al., 2010). The process could include a variable enhancement of the signal-to-noise ratio in the image (Peng et al., 2014).

Reconstructions were manually corrected and curated using a range of tools (e.g., virtual finger, polyline) in the Mozak extension (Zoran Popovic, Center for Game Science, University of Washington) of Terafly tools (Peng et al., 2014; Bria et al., 2016) in Vaa3D. Every attempt was made to generate a completely connected neuronal structure while remaining faithful to image data. If axonal processes could not be traced back to the main structure of the neuron, they were left unconnected. As a final step in the manual correction and curation process, an alternative analyst checked for missed branches or inappropriate connections. Once the reconstruction was deemed complete, multiple plugins were used to prepare neurons (saved as SWC files) for morphological analyses.

---

**Figure 1 – Figure supplement 1.**
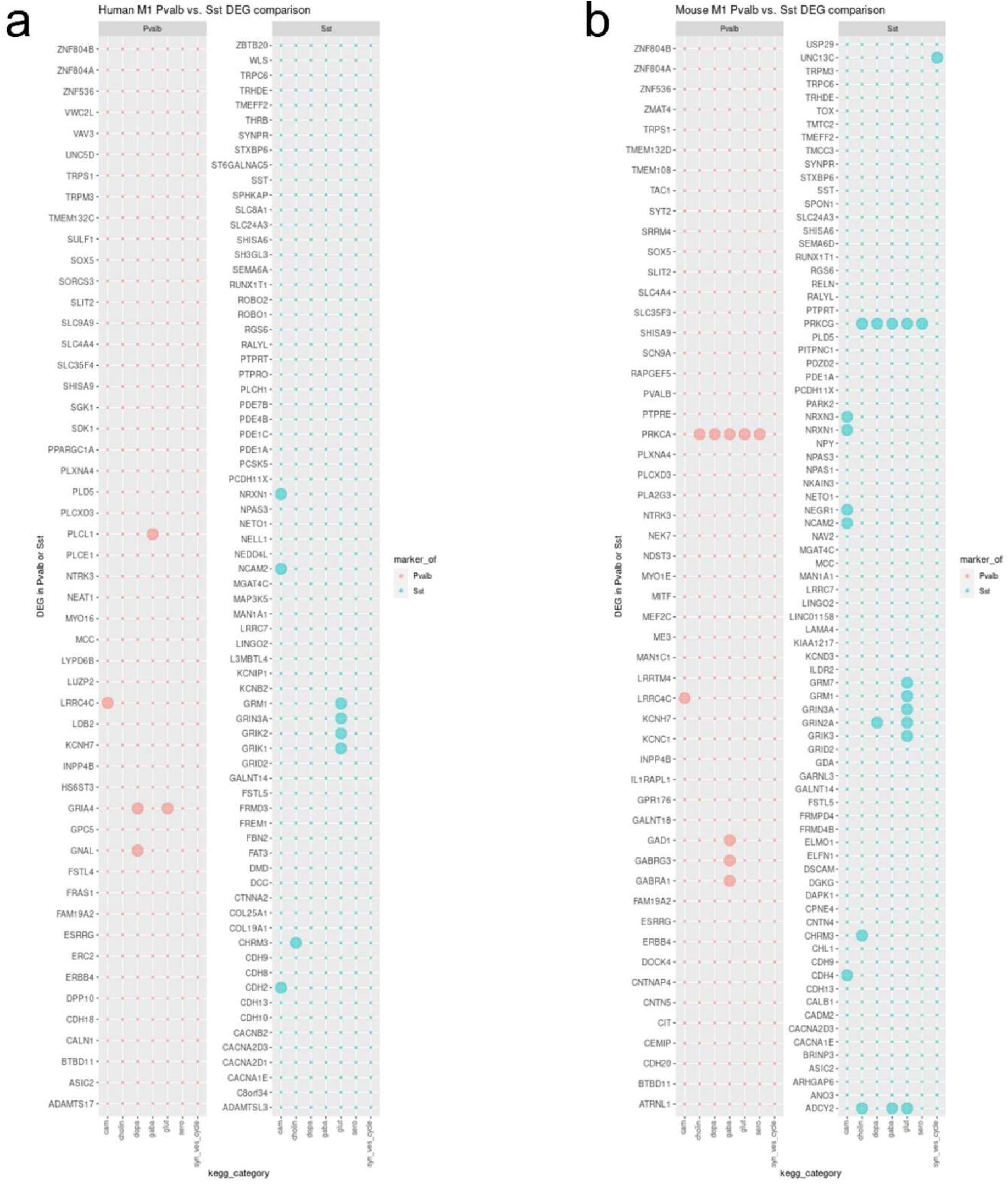
Differentially expressed genes (DEG) comparison by dot plot of Pvalb enriched DEG (red) and Sst enriched DEG (blue) in human (a) and mouse M1 datasets (b). Bigger dots show which KEGG pathway category they are associated with on the bottom. The bottom category are cell adhesion molecules (cam), synaptic gene associated with cholinergic (cholin), dopaminergic (dopa), GABAergic (gaba), glutamatergic (glut), serotoninergic (sero), or synaptic vesicles (syn_ves_cycle).

**Figure 2 – Figure supplement 1.**
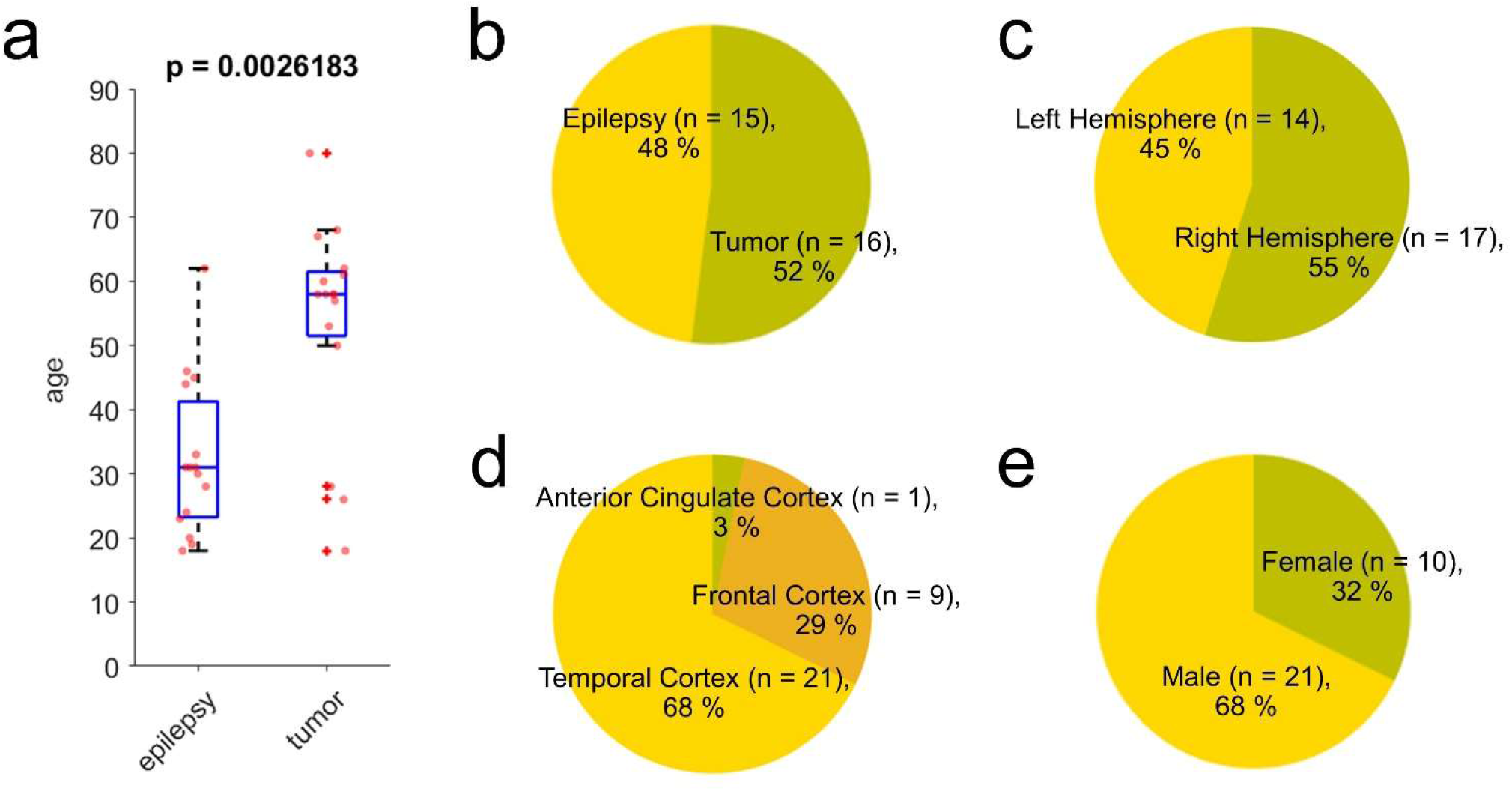
Summary of donor information. 31 donors were used for this study, including samples from epilepsy treatment and tumor removal surgeries. Patient ages were distributed from 18 to 80 and epilepsy patients were significantly younger than tumor patients (**a-b**). Brain areas, hemisphere, and sex distributions are displayed in **c-e**. P-value in **a** from Wilcoxon rank sum test.

**Figure 4 – Figure supplement 1.**
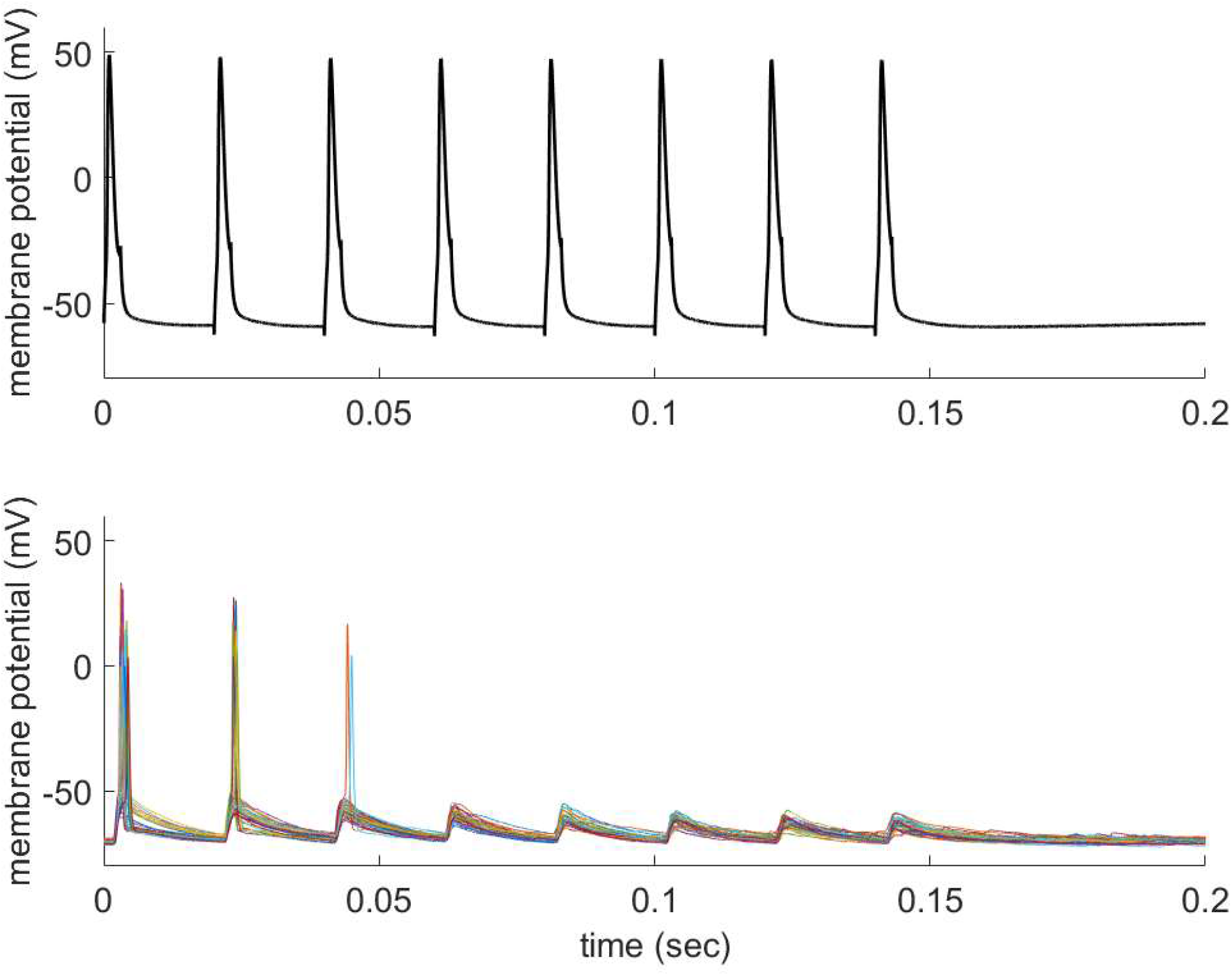
Example of presynaptic (pyramidal) unitary action potential and evoked postsynaptic (AAV labeled GABAergic) spike generation in organotypic slice culture when the postsynaptic cell was held at around −70 mV. Trains of 50 Hz presynaptic action potentials were repetitively generated by brief current injections (upper trace, 35 traces were averaged). Individual postsynaptic responses were overlayed with different colors (lower traces). Note that unitary EPSP evoked action potentials in some of trials especially in first 3 responses.

**Figure 4 – Figure supplement 2.**
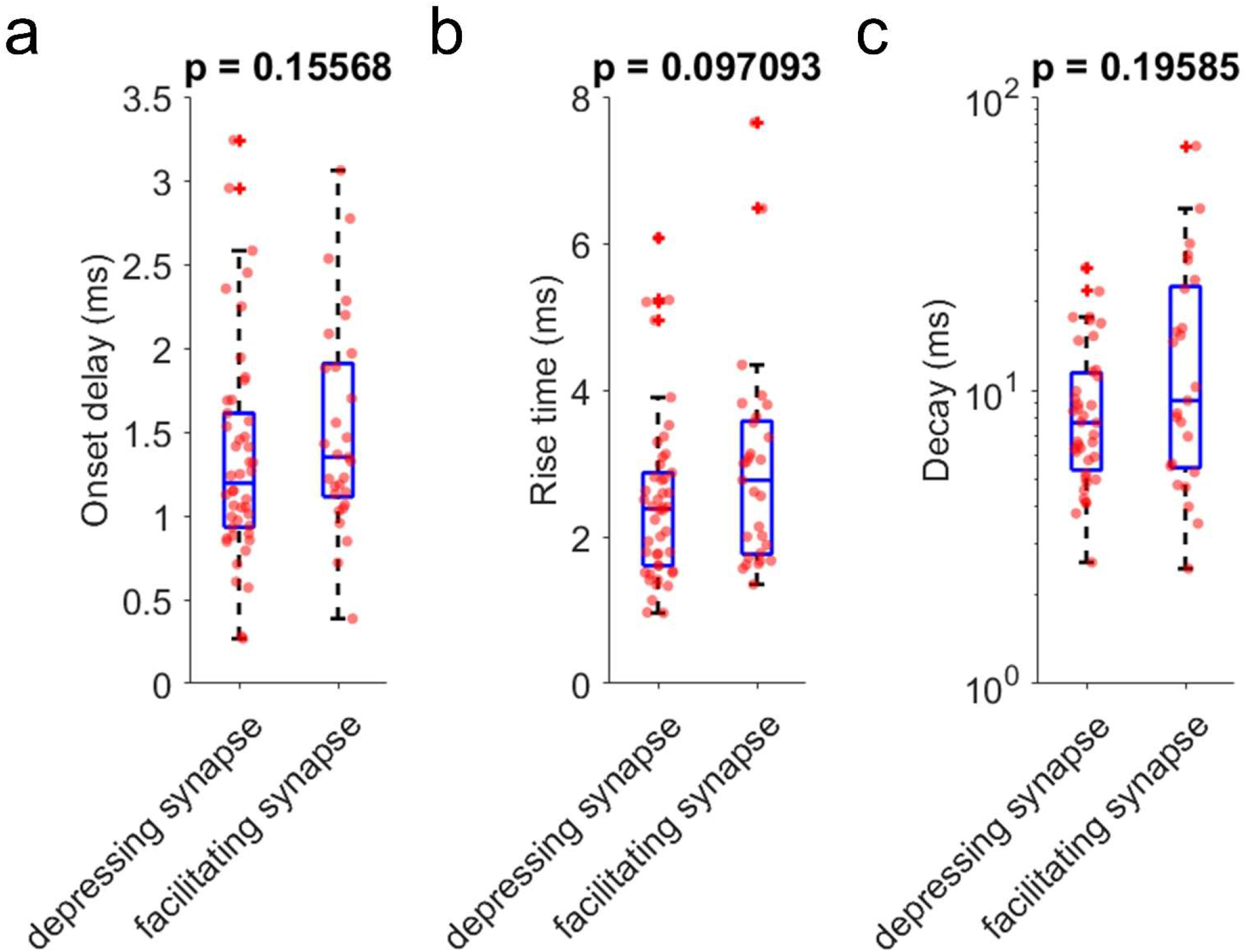
Comparison of EPSP kinetics based on their synaptic properties (i.e., depression and facilitation). Initial EPSP responses from 50 Hz stimulation were aligned from response onset and averaged (Figure 4i). 1:2 ratio was determinant for classifying synapses as depressing and facilitating at 50 Hz presynaptic stimulation (depressing synapses (*n* = 50) and facilitating synapses (*n* = 39)). **a**, Onset delay (*n* = 46, depressing synapses; *n* = 29, facilitating synapses). **b**, Rise time defined by time from response onset to peak (*n* = 46, depressing synapses; *n* = 29, facilitating synapses). **c**, Decay time constant (*n* = 39, depressing synapses; *n* = 25, facilitating synapses). P-values are from Wilcoxon rank sum test.

**Figure 4 – Figure supplement 3.**
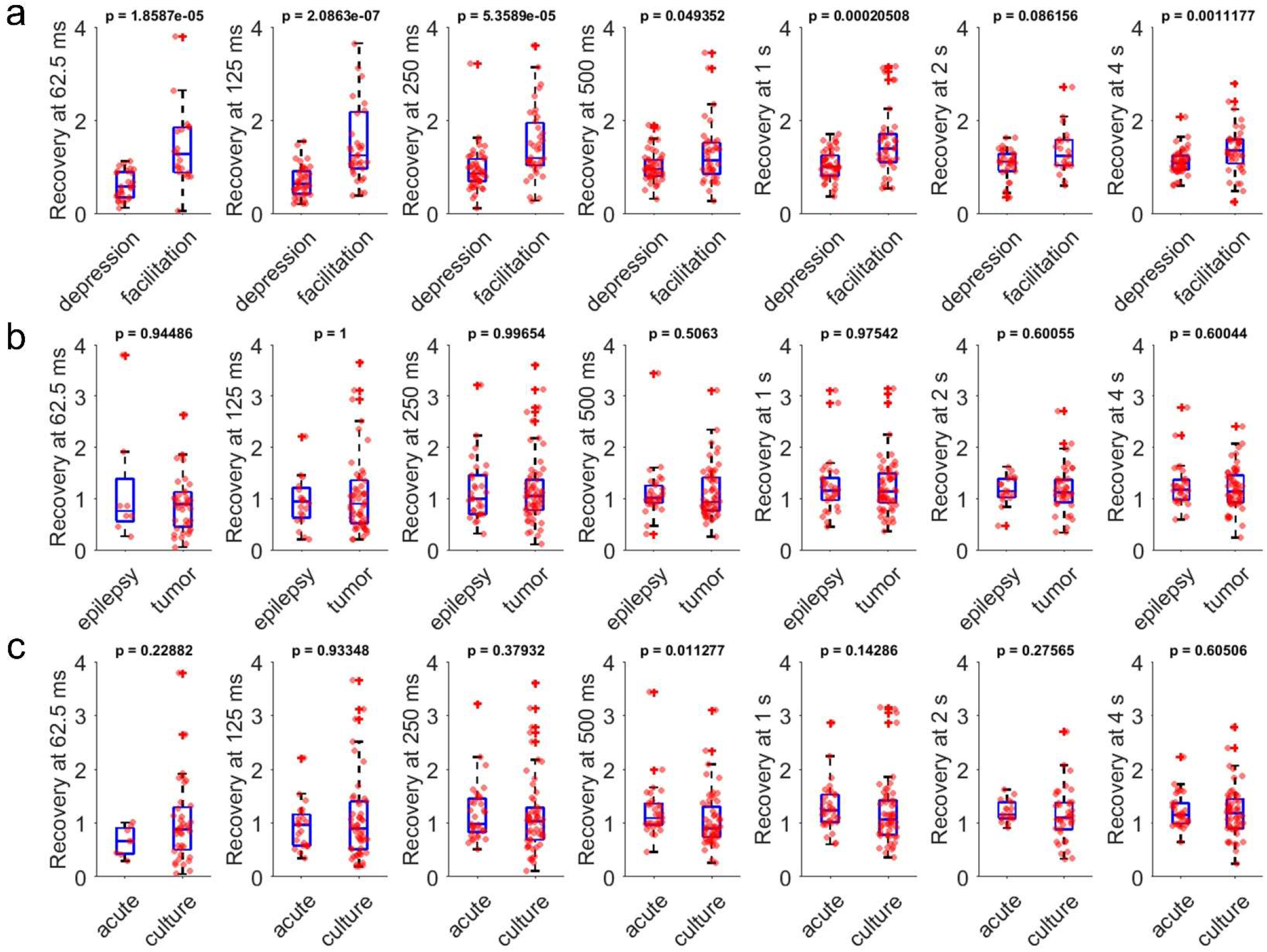
Comparison of recovery dynamics (at 62.5 ms, 125 ms, 250 ms, 500 ms, 1 s, 2s, 4 s) based on their synaptic properties (i.e., depression and facilitation), tissue origins (i.e., tumor or epilepsy) and preparation types (i.e., acute or slice culture). **a**, Normalized recovery (i.e., 9^th^ pulse responses divided by their first pulse responses) at various time delay (62.5, 125, 250, 500 ms, and 1, 2, 4 s) were compared between depressing (*n* = 50) and facilitating (*n* = 39) synapses defined by their 1:2 ratio at 50 Hz stimulation. **b**, Normalized recovery rates were compared from their tissue origins (*n* = 30, epilepsy; *n* = 59, tumor). **c,** Normalized recovery rates were compared from their tissue preparation types (*n* = 33, acute slice; *n* = 56, slice culture). P-values are from Wilcoxon rank sum test.

**Figure 4 – Figure supplement 4.**
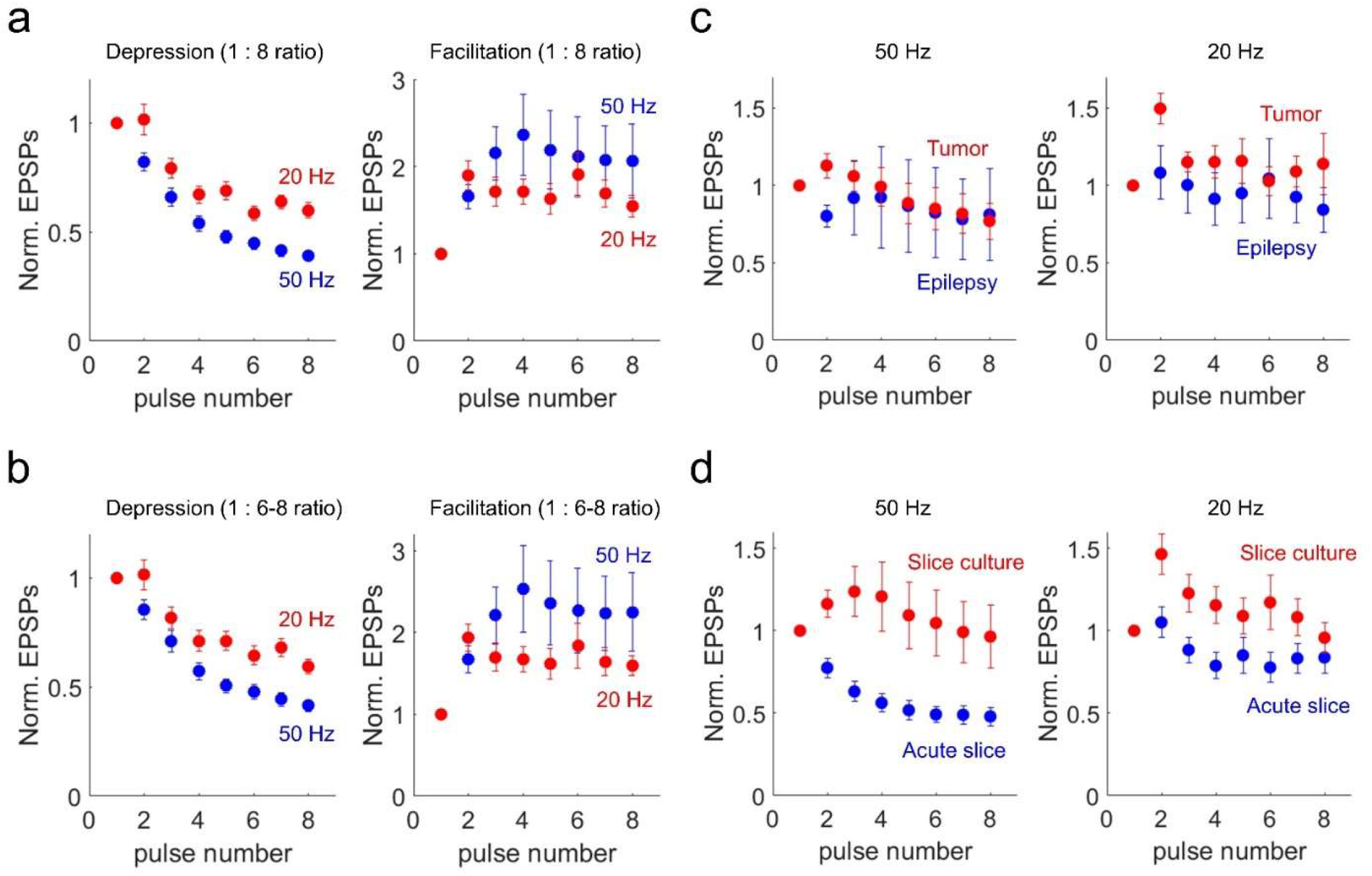
Presynaptic stimulation frequency dependent synaptic short-term dynamics with various synapse type definitions and comparison of synaptic short-term dynamics (50 Hz and 20 Hz) based on their tissue origins (i.e., tumor or epilepsy) and preparation types (i.e., acute or slice culture). **a,** Kinetics of synaptic dynamics. Depression and facilitation synapses were defined based on 1:8 ratio. Kinetics of dynamics from depressing synapses are displayed (*n* = 71, depression at 50 Hz, blue; *n* = 53, depression at 20 Hz, red; left). Similarly, kinetics of dynamics from facilitating synapses are displayed (*n* = 18, facilitation at 50 Hz; *n* = 26, facilitation at 20 Hz; right). **b,** Kinetics of synaptic dynamics. Depressing and facilitating synapses were defined based on 1: 6-8 ratio (average of 6 to 8 pulses compared to first pulse). Kinetics of dynamics from depressing synapses are displayed (*n* = 68, depression at 50 Hz, blue; *n* = 52, depression at 20 Hz, red; left). Similarly, kinetics of dynamics from facilitating synapses are displayed (*n* = 21, facilitation at 50 Hz; *n* = 27, facilitation at 20 Hz; right). Displayed data indicate mean ± s.e.m (error bars). **c**, Kinetics of synaptic dynamics were compared from their tissue origins. Left panel shows at 50 Hz (*n* = 30, epilepsy, blue; *n* = 59, tumor, red) and right panel shows at 20 Hz (*n* = 29, epilepsy, blue; *n* = 50, tumor, red). **d**, Kinetics of synaptic dynamics were compared from their slice preparations. Left panel shows at 50 Hz (*n* = 33, acute, blue; *n* = 56, slice culture, red) and note that only 4 out of 33 synapses showed synaptic facilitation at 50 Hz stimulation (1:2 ratio) in acute preparation and averaged values show strong depression. Right panel shows at 20 Hz (*n* = 29, acute slice, blue; *n* = 50, slice culture, red). Displayed data indicate mean ± s.e.m (error bars).

**Figure 5 – Figure supplement 1.**
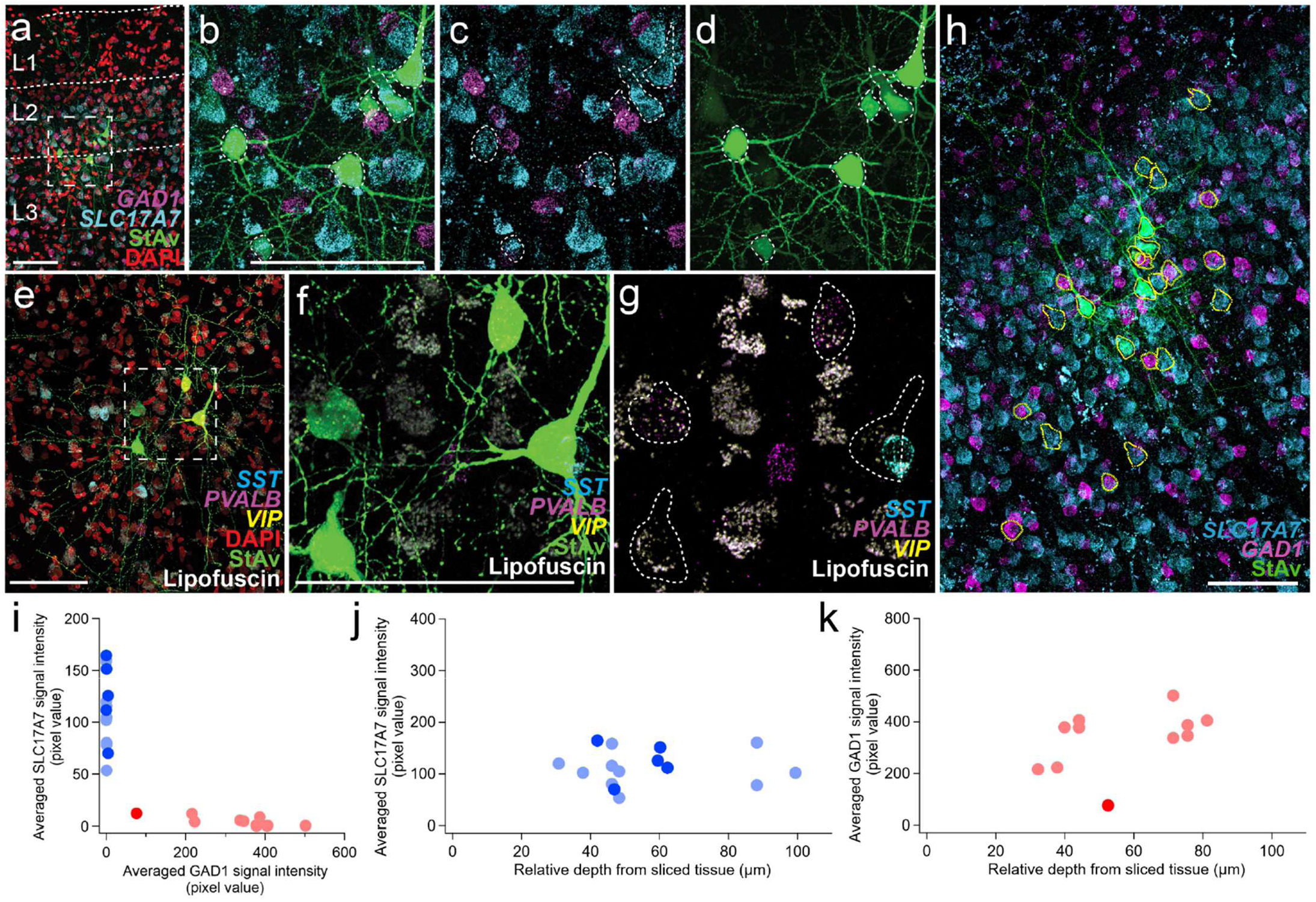
Two examples of MPC experiments with excitatory and inhibitory HCR markers in acute *ex vivo* human neocortex and depth dependence of HCR signals. Two examples MPC experiments shown in **a-d,h-k** and in **e-g**. **a,** Fluorescence montage of maximum intensity projections showing mFISH staining for excitatory (*SLC17A7*) and inhibitory (*GAD1*) neuron marker transcripts, nuclei (DAPI), and patched neurons (biocytin). Note lack of *SLC17A7* labeling in layer 1. (**b-d**) Enlargement of boxed region from **a. e**, Fluorescence montage showing four adjacent multi-patched cells from a different experiment. **f-g,** Two of the four patched neurons were labeled with *PVALB* transcript. Note, although the cell on the right overlaps with *SST*, this is a different Z-plane than the patched cells. No *VIP*-stained neurons were present in this field of view but were found outside of this region (data not shown). (**h-k**) Quantification of excitatory and inhibitory neuron markers in patched and unpatched neurons. **h**, Co-labeling for *SLC17A7* and *GAD1* mRNA. Yellow lines show 3D manually segmented neurons in a maximum intensity projection montage image. **i**, *SLC17A7* and *GAD1* expression is mutually exclusive. Blue indicates *SLC17A7^+^* neurons (patched neurons, blue; unpatched neurons, light blue) and red indicates *GAD1^+^* neurons (patched neurons, red; non-patched neurons, light red). The same cells are displayed as a function of cortical depth in **j-k**. **j**, *SLC17A7^+^* positive neurons are displayed along relative depth of the slice. **k**, Similarly, *GAD1^+^* positive neurons are displayed along relative depth of the slice. Most labeled cells showed similar mRNA intensity in patched and unpatched neurons, with the exception of one GAD1-positive neuron with lower transcript detection. Scale bars, 100 µm in **a,b,e,f,h**.

**Figure 5 – Figure supplement 2.**
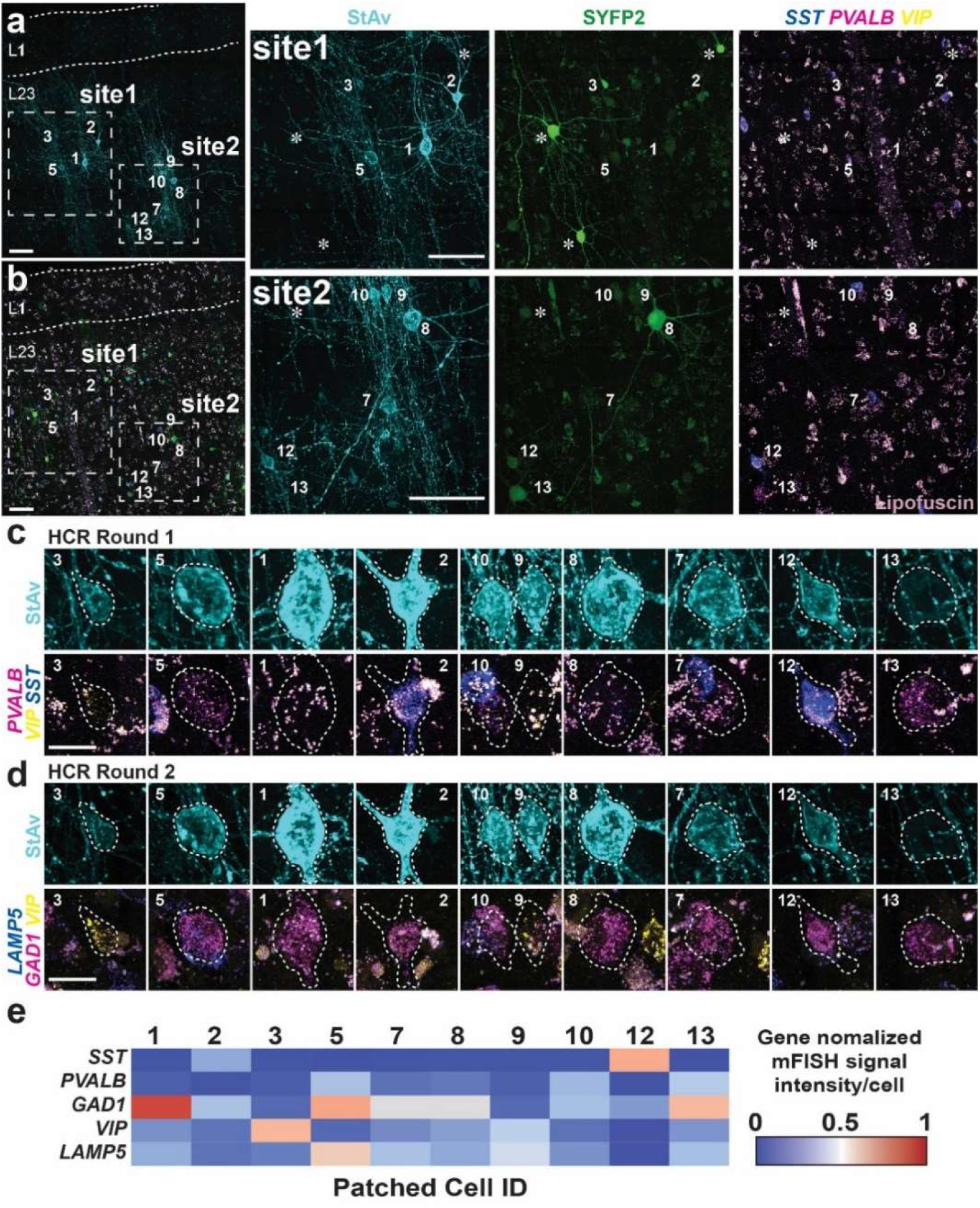
Examples of MPC experiments with inhibitory interneuron-targeted HCR staining. **a,** Fluorescent montage images of two neuron clusters evaluated by MPC recordings highlighted with boxes **site1** and **site2** with patched cell numbers after seven days in culture (7 DIV refer to 7 days *in vitro*). Patched neurons displayed with biocytin/streptavidin staining. **b,** Fluorescent montage of SYFP2 expression from a GABAergic neuron enhancer AAV (AAV-DLX2.0-SYFP2). **right panel,** high magnification images of **site1** and **site2** shown in **a** and **b**. Patched neurons displayed with biocytin/streptavidin only staining (StAv; left) or SYFP2 (SYFP2; middle), mFISH, and lipofuscin (light purple; right). Asterisks mark SYFP2^+^ cells not marked by biocytin/streptavidin. Scale bar, 100 µm. **c-d,** High resolution images of individual patched cells stained by mFISH in round 1 against *PVALB, SST*, and *VIP*, and round 2 against *LAMP5*, *GAD1*, and *VIP*. Scale bar, 10 µm. Cell numbers are labeled consistently in **a-b**. **e**, Expression level of each gene for each patched cell was quantified based on average intensity per cell. The average intensity was normalized by maximum value detected among the manually segmented patched non-patched cells shown in **Figure 5 – Figure supplement 3a**.

**Figure 5 – Figure supplement 3.**
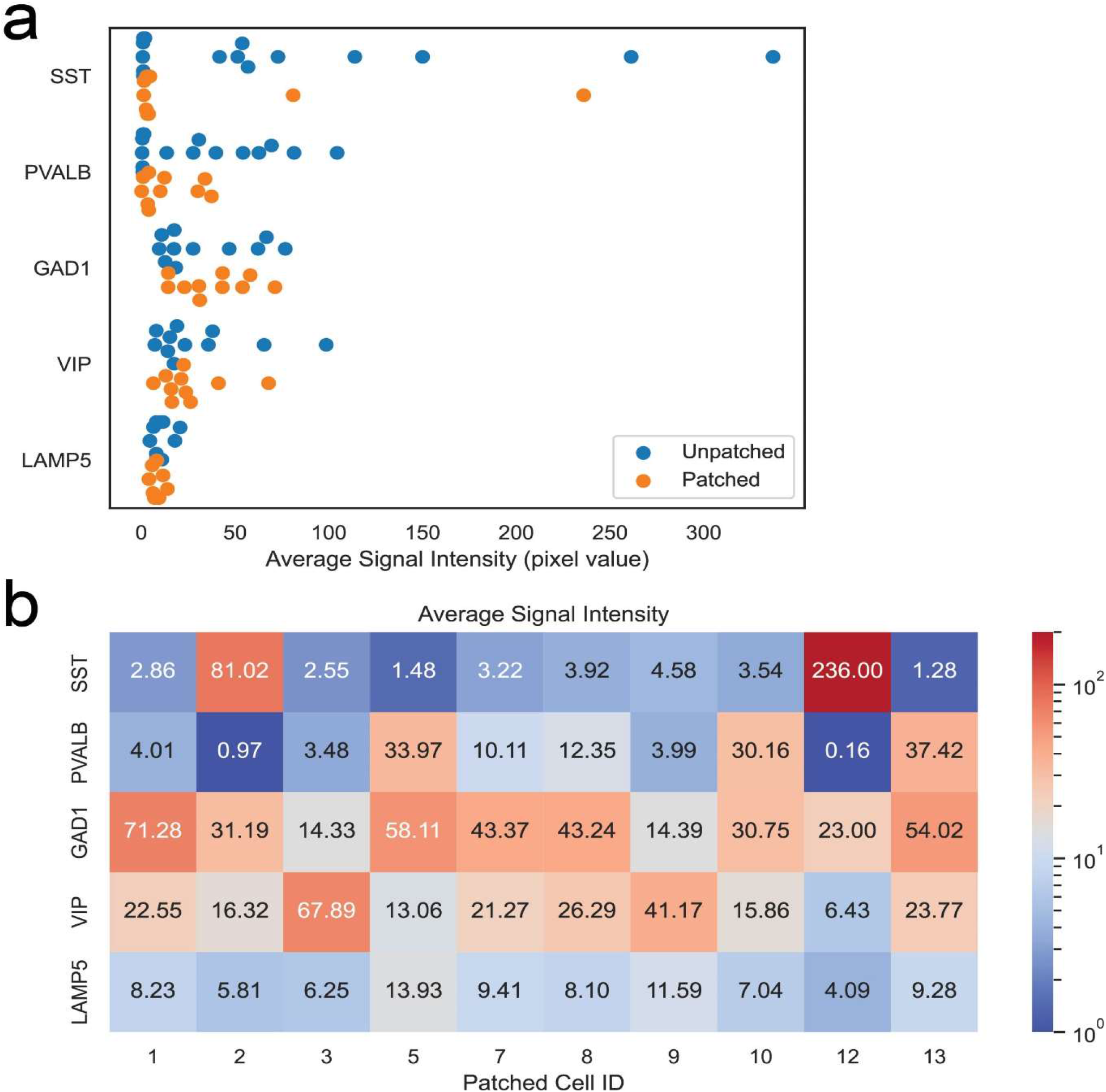
Quantification of HCR signal in slice culture preparation. **a**, HCR signal comparison between patched and non-patched neighboring neurons. Both patched and non-patched neighboring neurons were manually segmented in 3D. Average intensity values of HCR signals in each neuron were quantified and displayed. Orange closed circles indicate the patched neurons shown in Figure 3 and blue closed circles indicate non-patched neighboring neurons. **b**, Heat map displaying average fluorescence signal intensity values from manually segmented patched neurons. Patched cell numbers (Patched Cell ID) refer to the cells in Figure 5 **– Figure supplement 2e**.

**Figure 6 – Figure supplement 1.**
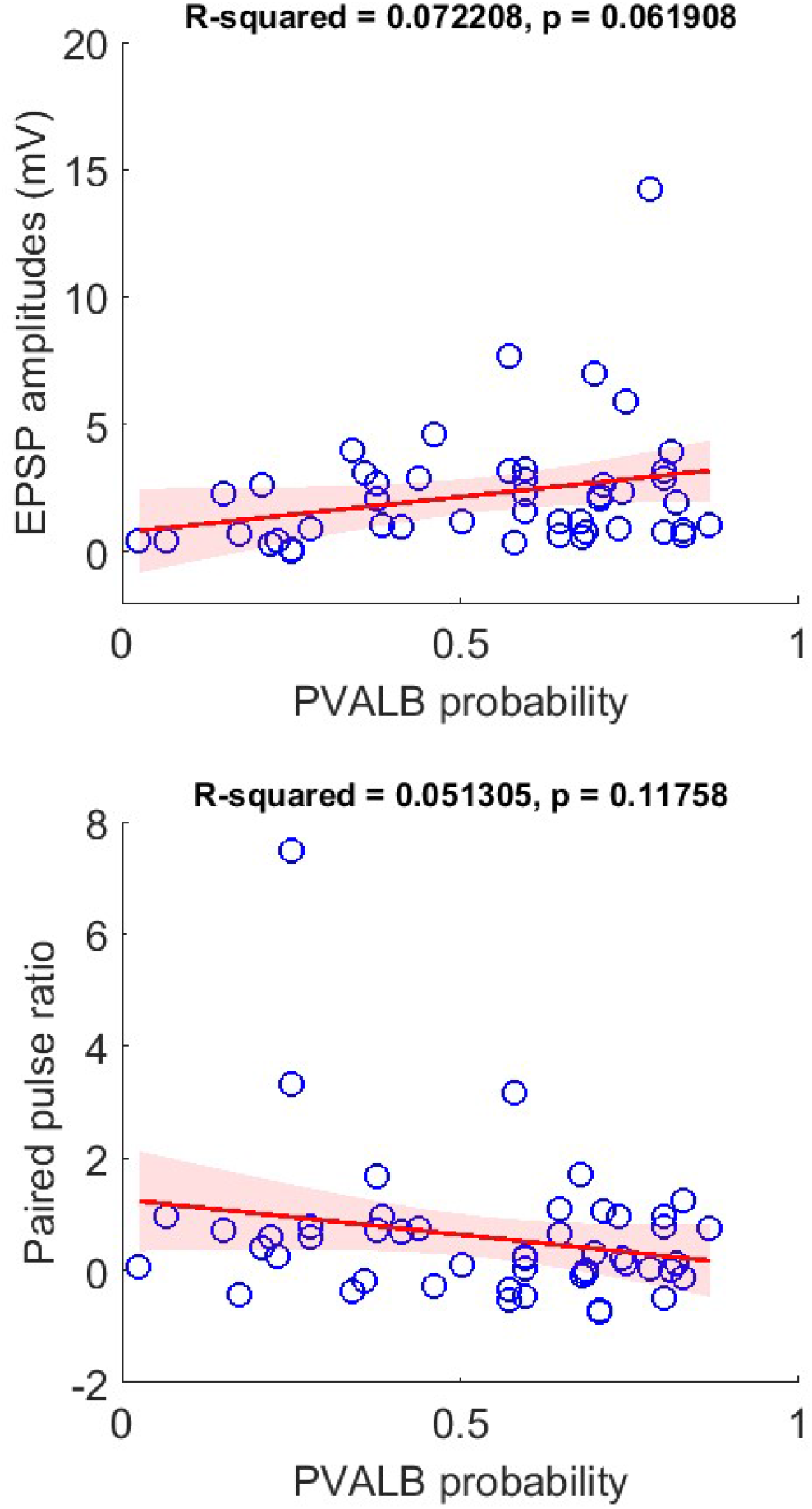
Classifier prediction with 20 Hz pulse trains. Lower correlation is shown between PVALB probability and their EPSP amplitudes (upper) or paired pulse (lower) with 20 Hz presynaptic stimulation protocol than 50 Hz stimuli. Regression line (red) with fitting confidence bounds (shaded region).

## References

1. Abbott, L. F., and Wade G. Regehr. 2004. “Synaptic Computation.” Nature 431 (7010): 796–803.

2. Andersson, My, Natalia Avaliani, Andreas Svensson, Jenny Wickham, Lars H. Pinborg, Bo Jespersen, Søren H. Christiansen, Johan Bengzon, David P. D. Woldbye, and Merab Kokaia. 2016. “Optogenetic Control of Human Neurons in Organotypic Brain Cultures.” Scientific Reports 6 (April): 24818.

3. Augustin, I., S. Korte, M. Rickmann, H. A. Kretzschmar, T. C. Südhof, J. W. Herms, and N. Brose. 2001. “The Cerebellum-Specific Munc13 Isoform Munc13-3 Regulates Cerebellar Synaptic Transmission and Motor Learning in Mice.” The Journal of Neuroscience: The Official Journal of the Society for Neuroscience 21 (1): 10–17.

4. Bakken, Trygve E., Nikolas L. Jorstad, Qiwen Hu, Blue B. Lake, Wei Tian, Brian E. Kalmbach, Megan Crow, et al. 2021. “Comparative Cellular Analysis of Motor Cortex in Human, Marmoset and Mouse.” Nature 598 (7879): 111–19.

5. Bankhead, Peter, Maurice B. Loughrey, José A. Fernández, Yvonne Dombrowski, Darragh G. McArt, Philip D. Dunne, Stephen McQuaid, et al. 2017. “QuPath: Open Source Software for Digital Pathology Image Analysis.” Scientific Reports 7 (1): 16878.

6. Bao, Jin, Kerstin Reim, and Takeshi Sakaba. 2010. “Target-Dependent Feedforward Inhibition Mediated by Short-Term Synaptic Plasticity in the Cerebellum.” The Journal of Neuroscience: The Official Journal of the Society for Neuroscience 30 (24): 8171–79.

7. Beaulieu-Laroche, Lou, Norma J. Brown, Marissa Hansen, Enrique H. S. Toloza, Jitendra Sharma, Ziv M. Williams, Matthew P. Frosch, Garth Rees Cosgrove, Sydney S. Cash, and Mark T. Harnett. 2021. “Allometric Rules for Mammalian Cortical Layer 5 Neuron Biophysics.” Nature 600 (7888): 274–78.

8. Beaulieu-Laroche, Lou, Enrique H. S. Toloza, Marie-Sophie van der Goes, Mathieu Lafourcade, Derrick Barnagian, Ziv M. Williams, Emad N. Eskandar, Matthew P. Frosch, Sydney S. Cash, and Mark T. Harnett. 2018. “Enhanced Dendritic Compartmentalization in Human Cortical Neurons.” Cell 175 (3): 643–51.e14.

9. Becht, Etienne, Leland McInnes, John Healy, Charles-Antoine Dutertre, Immanuel W. H. Kwok, Lai Guan Ng, Florent Ginhoux, and Evan W. Newell. 2018. “Dimensionality Reduction for Visualizing Single-Cell Data Using UMAP.” *Nature Biotechnology*, December. https://doi.org/10.1038/nbt.4314.

10. Beierlein, Michael, Jay R. Gibson, and Barry W. Connors. 2003. “Two Dynamically Distinct Inhibitory Networks in Layer 4 of the Neocortex.” Journal of Neurophysiology 90 (5): 2987–3000.

11. Berger, Thomas K., Rodrigo Perin, Gilad Silberberg, and Henry Markram. 2009. “Frequency-Dependent Disynaptic Inhibition in the Pyramidal Network: A Ubiquitous Pathway in the Developing Rat Neocortex.” The Journal of Physiology 587 (Pt 22): 5411–25.

12. Berg, Jim, Staci A. Sorensen, Jonathan T. Ting, Jeremy A. Miller, Thomas Chartrand, Anatoly Buchin, Trygve E. Bakken, et al. 2021. “Human Neocortical Expansion Involves Glutamatergic Neuron Diversification.” Nature 598 (7879): 151–58.

13. Blackman, Arne V., Therese Abrahamsson, Rui Ponte Costa, Txomin Lalanne, and P. Jesper Sjöström. 2013. “Target-Cell-Specific Short-Term Plasticity in Local Circuits.” Frontiers in Synaptic Neuroscience 5 (December): 11.

14. Boldog, Eszter, Trygve E. Bakken, Rebecca D. Hodge, Mark Novotny, Brian D. Aevermann, Judith Baka, Sándor Bordé, et al. 2018. “Transcriptomic and Morphophysiological Evidence for a Specialized Human Cortical GABAergic Cell Type.” Nature Neuroscience 21 (9): 1185–95.

15. Bria, Alessandro, Giulio Iannello, Leonardo Onofri, and Hanchuan Peng. 2016. “TeraFly: Real-Time Three-Dimensional Visualization and Annotation of Terabytes of Multidimensional Volumetric Images.” Nature Methods 13 (3): 192–94.

16. Bugeon, Stephane, Joshua Duffield, Mario Dipoppa, Anne Ritoux, Isabelle Prankerd, Dimitris Nicolout-sopoulos, David Orme, et al. 2021. “A Transcriptomic Axis Predicts State Modulation of Cortical Interneurons.” bioRxiv. https://doi.org/10.1101/2021.10.24.465600.

17. Cadwell, Cathryn R., Athanasia Palasantza, Xiaolong Jiang, Philipp Berens, Qiaolin Deng, Marlene Yilmaz, Jacob Reimer, et al. 2016. “Electrophysiological, Transcriptomic and Morphologic Profiling of Single Neurons Using Patch-Seq.” Nature Biotechnology 34 (2): 199–203.

18. Campagnola, Luke, Megan B. Kratz, and Paul B. Manis. 2014. “ACQ4: An Open-Source Software Platform for Data Acquisition and Analysis in Neurophysiology Research.” Frontiers in Neuroinformatics 8 (January): 3.

19. Campagnola, Luke, Stephanie C. Seeman, Thomas Chartrand, Lisa Kim, Alex Hoggarth, Clare Gamlin, Shinya Ito, et al. 2021. “Local Connectivity and Synaptic Dynamics in Mouse and Human Neocortex.” bioRxiv. bioRxiv. https://doi.org/10.1101/2021.03.31.437553.

20. Chan, Ken Y., Min J. Jang, Bryan B. Yoo, Alon Greenbaum, Namita Ravi, Wei-Li Wu, Luis Sánchez-Guardado, et al. 2017. “Engineered AAVs for Efficient Noninvasive Gene Delivery to the Central and Peripheral Nervous Systems.” Nature Neuroscience 20 (8): 1172–79.

21. Chen, Kok Hao, Alistair N. Boettiger, Jeffrey R. Moffitt, Siyuan Wang, and Xiaowei Zhuang. 2015. “RNA Imaging. Spatially Resolved, Highly Multiplexed RNA Profiling in Single Cells.” Science 348 (6233): aaa6090.

22. Chen, Shuo, Yuzhou Chang, Liangping Li, Diana Acosta, Cody Morrison, Cankun Wang, Dominic Julian, et al. 2021. “Spatially Resolved Transcriptomics Reveals Unique Gene Signatures Associated with Human Temporal Cortical Architecture and Alzheimer’s Pathology.” bioRxiv. https://doi.org/10.1101/2021.07.07.451554.

23. Chittajallu, Ramesh, Kurt Auville, Vivek Mahadevan, Mandy Lai, Steven Hunt, Daniela Calvigioni, Kenneth A. Pelkey, Kareem A. Zaghloul, and Chris J. McBain. 2020. “Activity-Dependent Tuning of Intrinsic Excitability in Mouse and Human Neurogliaform Cells.” eLife 9 (June). https://doi.org/10.7554/eLife.57571.

24. Choi, Harry M. T., Victor A. Beck, and Niles A. Pierce. 2014. “Next-Generation in Situ Hybridization Chain Reaction: Higher Gain, Lower Cost, Greater Durability.” ACS Nano 8 (5): 4284–94.

25. Choi, Harry M. T., Joann Y. Chang, Le A. Trinh, Jennifer E. Padilla, Scott E. Fraser, and Niles A. Pierce. 2010. “Programmable in Situ Amplification for Multiplexed Imaging of mRNA Expression.” Nature Biotechnology 28 (11): 1208–12.

26. Choi, Harry M. T., Maayan Schwarzkopf, Mark E. Fornace, Aneesh Acharya, Georgios Artavanis, Johannes Stegmaier, Alexandre Cunha, and Niles A. Pierce. 2018. “Third-Generation Hybridization Chain Reaction: Multiplexed, Quantitative, Sensitive, Versatile, Robust.” Development 145 (12). https://doi.org/10.1242/dev.165753.

27. Condylis, Cameron, Abed Ghanbari, Nikita Manjrekar, Karina Bistrong, Shenqin Yao, Zizhen Yao, Thuc Nghi Nguyen, Hongkui Zeng, Bosiljka Tasic, and Jerry L. Chen. 2022. “Dense Functional and Molecular Readout of a Circuit Hub in Sensory Cortex.” Science 375 (6576): eabl5981.

28. Daigle, Tanya L., Linda Madisen, Travis A. Hage, Matthew T. Valley, Ulf Knoblich, Rylan S. Larsen, Marc M. Takeno, et al. 2018. “A Suite of Transgenic Driver and Reporter Mouse Lines with Enhanced Brain-Cell-Type Targeting and Functionality.” Cell 174 (2): 465–80.e22.

29. Dimidschstein, Jordane, Qian Chen, Robin Tremblay, Stephanie L. Rogers, Giuseppe-Antonio Saldi, Lihua Guo, Qing Xu, et al. 2016. “A Viral Strategy for Targeting and Manipulating Interneurons across Vertebrate Species.” Nature Neuroscience 19 (12): 1743–49.

30. Eng, Chee-Huat Linus, Michael Lawson, Qian Zhu, Ruben Dries, Noushin Koulena, Yodai Takei, Jina Yun, et al. 2019. “Transcriptome-Scale Super-Resolved Imaging in Tissues by RNA seqFISH.” Nature 568 (7751): 235–39.

31. Eugène, Emmanuel, Françoise Cluzeaud, Carmen Cifuentes-Diaz, Desdemona Fricker, Caroline Le Duigou, Stephane Clemenceau, Michel Baulac, Jean-Christophe Poncer, and Richard Miles. 2014. “An Organotypic Brain Slice Preparation from Adult Patients with Temporal Lobe Epilepsy.” Journal of Neuroscience Methods 235 (September): 234–44.

32. Futai, Kensuke, Myung Jong Kim, Tsutomu Hashikawa, Peter Scheiffele, Morgan Sheng, and Yasunori Hayashi. 2007. “Retrograde Modulation of Presynaptic Release Probability through Signaling Mediated by PSD-95–neuroligin.” Nature Neuroscience. https://doi.org/10.1038/nn1837.

33. Fu, Yu, Jason M. Tucciarone, J. Sebastian Espinosa, Nengyin Sheng, Daniel P. Darcy, Roger A. Nicoll, Z. Josh Huang, and Michael P. Stryker. 2014. “A Cortical Circuit for Gain Control by Behavioral State.” Cell 156 (6): 1139–52.

34. Gouwens, Nathan W., Staci A. Sorensen, Jim Berg, Changkyu Lee, Tim Jarsky, Jonathan Ting, Susan M. Sunkin, et al. 2019. “Classification of Electrophysiological and Morphological Neuron Types in the Mouse Visual Cortex.” Nature Neuroscience 22 (7): 1182–95.

35. Graybuck, Lucas T., Tanya L. Daigle, Adriana E. Sedeño-Cortés, Miranda Walker, Brian Kalmbach, Garreck H. Lenz, Thuc Nghi Nguyen, et al. n.d. “Enhancer Viruses and a Transgenic Platform for Combinatorial Cell Subclass-Specific Labeling.” https://doi.org/10.1101/525014.

36. Hao, Yuhan, Stephanie Hao, Erica Andersen-Nissen, William M. Mauck 3rd, Shiwei Zheng, Andrew Butler, Maddie J. Lee, et al. 2021. “Integrated Analysis of Multimodal Single-Cell Data.” Cell 184 (13): 3573–87.e29.

37. He, Miao, Jason Tucciarone, Soohyun Lee, Maximiliano José Nigro, Yongsoo Kim, Jesse Maurica Levine, Sean Michael Kelly, et al. 2016. “Strategies and Tools for Combinatorial Targeting of GABAergic Neurons in Mouse Cerebral Cortex.” Neuron 92 (2): 555.

38. Hill, Robert Sean, and Christopher A. Walsh. 2005. “Molecular Insights into Human Brain Evolution.” Nature 437 (7055): 64–67.

39. Hodge, Rebecca D., Trygve E. Bakken, Jeremy A. Miller, Kimberly A. Smith, Eliza R. Barkan, Lucas T. Graybuck, Jennie L. Close, et al. 2019. “Conserved Cell Types with Divergent Features in Human versus Mouse Cortex.” Nature 573 (7772): 61–68.

40. Hrvatin, Sinisa, Christopher P. Tzeng, M. Aurel Nagy, Hume Stroud, Charalampia Koutsioumpa, Oren F. Wilcox, Elena G. Assad, et al. 2019. “A Scalable Platform for the Development of Cell-Type-Specific Viral Drivers.” eLife 8 (September). https://doi.org/10.7554/eLife.48089.

41. Huang, Z. Josh, and Anirban Paul. 2019. “The Diversity of GABAergic Neurons and Neural Communication Elements.” Nature Reviews. Neuroscience 20 (9): 563–72.

42. Isaacson, Jeffry S., and Massimo Scanziani. 2011. “How Inhibition Shapes Cortical Activity.” Neuron 72 (2): 231–43.

43. Jiang, Man, Jie Zhu, Yaping Liu, Mingpo Yang, Cuiping Tian, Shan Jiang, Yonghong Wang, Hui Guo, Kaiyan Wang, and Yousheng Shu. 2012. “Enhancement of Asynchronous Release from Fast-Spiking Interneuron in Human and Rat Epileptic Neocortex.” PLoS Biology 10 (5): e1001324.

44. Jüttner, Josephine, Arnold Szabo, Brigitte Gross-Scherf, Rei K. Morikawa, Santiago B. Rompani, Peter Hantz, Tamas Szikra, et al. 2019. “Targeting Neuronal and Glial Cell Types with Synthetic Promoter AAVs in Mice, Non-Human Primates and Humans.” Nature Neuroscience 22 (8): 1345–56.

45. Kalmbach, Brian E., Anatoly Buchin, Brian Long, Jennie Close, Anirban Nandi, Jeremy A. Miller, Trygve E. Bakken, et al. 2018. “H-Channels Contribute to Divergent Intrinsic Membrane Properties of Supragranular Pyramidal Neurons in Human versus Mouse Cerebral Cortex.” Neuron. https://doi.org/10.1016/j.neuron.2018.10.012.

46. Koester, Helmut J., and Daniel Johnston. 2005. “Target Cell-Dependent Normalization of Transmitter Release at Neocortical Synapses.” Science 308 (5723): 863–66.

47. Komlósi, Gergely, Gábor Molnár, Márton Rózsa, Szabolcs Oláh, Pál Barzó, and Gábor Tamás. 2012. “Fluoxetine (prozac) and Serotonin Act on Excitatory Synaptic Transmission to Suppress Single Layer 2/3 Pyramidal Neuron-Triggered Cell Assemblies in the Human Prefrontal Cortex.” The Journal of Neuroscience: The Official Journal of the Society for Neuroscience 32 (46): 16369–78.

48. Lake, Blue B., Rizi Ai, Gwendolyn E. Kaeser, Neeraj S. Salathia, Yun C. Yung, Rui Liu, Andre Wildberg, et al. 2016. “Neuronal Subtypes and Diversity Revealed by Single-Nucleus RNA Sequencing of the Human Brain.” Science 352 (6293): 1586–90.

49. Le Duigou, Caroline, Etienne Savary, Mélanie Morin-Brureau, Daniel Gomez-Dominguez, André Sobczyk, Farah Chali, Giampaolo Milior, et al. 2018. “Imaging Pathological Activities of Human Brain Tissue in Organotypic Culture.” Journal of Neuroscience Methods 298 (March): 33–44.

50. Lee, Brian R., Agata Budzillo, Kristen Hadley, Jeremy A. Miller, Tim Jarsky, Katherine Baker, Dijon Hill, et al. 2021. “Scaled, High Fidelity Electrophysiological, Morphological, and Transcriptomic Cell Characterization.” eLife 10 (August). https://doi.org/10.7554/eLife.65482.

51. Lee, Soohyun, Illya Kruglikov, Z. Josh Huang, Gord Fishell, and Bernardo Rudy. 2013. “A Disinhibitory Circuit Mediates Motor Integration in the Somatosensory Cortex.” Nature Neuroscience. https://doi.org/10.1038/nn.3544.

52. Letzkus, Johannes J., Steffen B. E. Wolff, Elisabeth M. M. Meyer, Philip Tovote, Julien Courtin, Cyril Herry, and Andreas Lüthi. 2011. “A Disinhibitory Microcircuit for Associative Fear Learning in the Auditory Cortex.” Nature. https://doi.org/10.1038/nature10674.

53. Lukacs, Istvan Paul, Ruggiero Francavilla, Martin Field, Emily Hunter, Michael Howarth, Sawa Horie, Puneet Plaha, et al. 2022. “Differential Effects of Group III Metabotropic Glutamate Receptors on Spontaneous Inhibitory Synaptic Currents in Spine-Innervating Double Bouquet and Parvalbumin-Expressing Dendrite-Targeting GABAergic Interneurons in Human Neocortex.” bioRxiv. https://doi.org/10.1101/2022.03.05.483105.

54. Mansvelder, Huibert D., Matthijs B. Verhoog, and Natalia A. Goriounova. 2019. “Synaptic Plasticity in Human Cortical Circuits: Cellular Mechanisms of Learning and Memory in the Human Brain?” Current Opinion in Neurobiology 54 (February): 186–93.

55. Mehta, Preeti, Lauren Kreeger, Dennis C. Wylie, Jagruti J. Pattadkal, Tara Lusignan, Matthew J. Davis, Gergely F. Turi, et al. 2019. “Functional Access to Neuron Subclasses in Rodent and Primate Forebrain.” Cell Reports 26 (10): 2818–32.e8.

56. Mich, John K., Lucas T. Graybuck, Erik E. Hess, Joseph T. Mahoney, Yoshiko Kojima, Yi Ding, Saroja Somasundaram, et al. 2021. “Functional Enhancer Elements Drive Subclass-Selective Expression from Mouse to Primate Neocortex.” Cell Reports 34 (13): 108754.

57. Molnár, Gábor, Szabolcs Oláh, Gergely Komlósi, Miklós Füle, János Szabadics, Csaba Varga, Pál Barzó, and Gábor Tamás. 2008. “Complex Events Initiated by Individual Spikes in the Human Cerebral Cortex.” PLoS Biology 6 (9): e222.

58. Molnár, Gábor, Márton Rózsa, Judith Baka, Noémi Holderith, Pál Barzó, Zoltan Nusser, and Gábor Tamás. 2016. “Human Pyramidal to Interneuron Synapses Are Mediated by Multi-Vesicular Release and Multiple Docked Vesicles.” eLife 5 (August). https://doi.org/10.7554/eLife.18167.

59. Nair, Rajeevkumar Raveendran, Stefan Blankvoort, Maria Jose Lagartos, and Cliff Kentros. 2019. “Generation of Viral Vectors Specific to Neuronal Subtypes of Targeted Brain Regions by Enhancer-Driven Gene Expression (EDGE).” bioRxiv. bioRxiv. https://doi.org/10.1101/606467.

60. Obermayer, Joshua, Tim S. Heistek, Amber Kerkhofs, Natalia A. Goriounova, Tim Kroon, Johannes C. Baayen, Sander Idema, Guilherme Testa-Silva, Jonathan J. Couey, and Huibert D. Mansvelder. 2018. “Lateral Inhibition by Martinotti Interneurons Is Facilitated by Cholinergic Inputs in Human and Mouse Neocortex.” Nature Communications 9 (1): 4101.

61. Paul, Anirban, Megan Crow, Ricardo Raudales, Miao He, Jesse Gillis, and Z. Josh Huang. 2017. “Transcriptional Architecture of Synaptic Communication Delineates GABAergic Neuron Identity.” Cell 171 (3): 522–39.e20.

62. Peng, Hanchuan, Alessandro Bria, Zhi Zhou, Giulio Iannello, and Fuhui Long. 2014. “Extensible Visualization and Analysis for Multidimensional Images Using Vaa3D.” Nature Protocols 9 (1): 193– 208.

63. Peng, Hanchuan, Zongcai Ruan, Fuhui Long, Julie H. Simpson, and Eugene W. Myers. 2010. “V3D Enables Real-Time 3D Visualization and Quantitative Analysis of Large-Scale Biological Image Data Sets.” Nature Biotechnology 28 (4): 348–53.

64. Peng, Yangfan, Franz Xaver Mittermaier, Henrike Planert, Ulf Christoph Schneider, Henrik Alle, and Jörg Rolf Paul Geiger. 2019. “High-Throughput Microcircuit Analysis of Individual Human Brains through next-Generation Multineuron Patch-Clamp.” eLife 8 (November). https://doi.org/10.7554/eLife.48178.

65. Pfeffer, Carsten K., Mingshan Xue, Miao He, Z. Josh Huang, and Massimo Scanziani. 2013. “Inhibition of Inhibition in Visual Cortex: The Logic of Connections between Molecularly Distinct Interneurons.” Nature Neuroscience 16 (8): 1068–76.

66. Pi, Hyun-Jae, Balázs Hangya, Duda Kvitsiani, Joshua I. Sanders, Z. Josh Huang, and Adam Kepecs. 2013. “Cortical Interneurons That Specialize in Disinhibitory Control.” Nature. https://doi.org/10.1038/nature12676.

67. Planert, Henrike, Franz X. Mittermaier, Sabine Grosser, Pawel Fidzinski, Ulf C. Schneider, Helena Radbruch, Julia Onken, et al. 2021. “Intra-Individual Physiomic Landscape of Pyramidal Neurons in the Human Neocortex.” bioRxiv. https://doi.org/10.1101/2021.11.08.467668.

68. Poorthuis, Rogier B., Karzan Muhammad, Mantian Wang, Matthijs B. Verhoog, Stephan Junek, Anne Wrana, Huibert D. Mansvelder, and Johannes J. Letzkus. 2018. “Rapid Neuromodulation of Layer 1 Interneurons in Human Neocortex.” Cell Reports 23 (4): 951–58.

69. Pouille, Frédéric, and Massimo Scanziani. 2004. “Routing of Spike Series by Dynamic Circuits in the Hippocampus.” Nature 429 (6993): 717–23.

70. Reyes, A., R. Lujan, A. Rozov, N. Burnashev, P. Somogyi, and B. Sakmann. 1998. “Target-Cell-Specific Facilitation and Depression in Neocortical Circuits.” Nature Neuroscience 1 (4): 279–85.

71. Schwarz, Niklas, Ulrike B. S. Hedrich, Hannah Schwarz, Harshad P A, Nele Dammeier, Eva Auffenberg, Francesco Bedogni, et al. 2017. “Human Cerebrospinal Fluid Promotes Long-Term Neuronal Viability and Network Function in Human Neocortical Organotypic Brain Slice Cultures.” Scientific Reports 7 (1): 12249.

72. Schwarz, Niklas, Betül Uysal, Marc Welzer, Jacqueline C. Bahr, Nikolas Layer, Heidi Löffler, Kornelijus Stanaitis, et al. 2019. “Long-Term Adult Human Brain Slice Cultures as a Model System to Study Human CNS Circuitry and Disease.” eLife 8 (September). https://doi.org/10.7554/eLife.48417.

73. Seeman, Stephanie C., Luke Campagnola, Pasha A. Davoudian, Alex Hoggarth, Travis A. Hage, Alice Bosma-Moody, Christopher A. Baker, et al. 2018. “Sparse Recurrent Excitatory Connectivity in the Microcircuit of the Adult Mouse and Human Cortex.” eLife 7 (September). https://doi.org/10.7554/eLife.37349.

74. Shah, Sheel, Eric Lubeck, Maayan Schwarzkopf, Ting-Fang He, Alon Greenbaum, Chang Ho Sohn, Antti Lignell, et al. 2016. “Single-Molecule RNA Detection at Depth by Hybridization Chain Reaction and Tissue Hydrogel Embedding and Clearing.” Development 143 (15): 2862–67.

75. Silberberg, Gilad, and Henry Markram. 2007. “Disynaptic Inhibition between Neocortical Pyramidal Cells Mediated by Martinotti Cells.” Neuron 53 (5): 735–46.

76. Smith, Stephen J., Uygar Sümbül, Lucas T. Graybuck, Forrest Collman, Sharmishtaa Seshamani, Rohan Gala, Olga Gliko, et al. 2019. “Single-Cell Transcriptomic Evidence for Dense Intracortical Neuropeptide Networks.” eLife 8 (November). https://doi.org/10.7554/eLife.47889.

77. Stachniak, Tevye Jason, Emily Lauren Sylwestrak, Peter Scheiffele, Benjamin J. Hall, and Anirvan Ghosh. 2019. “Elfn1-Induced Constitutive Activation of mGluR7 Determines Frequency-Dependent Recruitment of Somatostatin Interneurons.” The Journal of Neuroscience: The Official Journal of the Society for Neuroscience 39 (23): 4461–74.

78. Stühmer, Thorsten, Luis Puelles, Marc Ekker, and John L. R. Rubenstein. 2002. “Expression from a Dlx Gene Enhancer Marks Adult Mouse Cortical GABAergic Neurons.” Cerebral Cortex 12 (1): 75–85.

79. Subkhankulova, Tatiana, Kojiro Yano, Hugh P. C. Robinson, and Frederick J. Livesey. 2010. “Grouping and Classifying Electrophysiologically-Defined Classes of Neocortical Neurons by Single Cell, Whole-Genome Expression Profiling.” Frontiers in Molecular Neuroscience 3 (April): 10.

80. Suriano, Christos M., Jessica L. Verpeut, Neerav Kumar, Jie Ma, Caroline Jung, and Lisa M. Boulanger. n.d. “Adeno-Associated Virus (AAV) Reduces Cortical Dendritic Complexity in a TLR9-Dependent Manner.” https://doi.org/10.1101/2021.09.28.462148.

81. Sylwestrak, Emily L., and Anirvan Ghosh. 2012. “Elfn1 Regulates Target-Specific Release Probability at CA1-Interneuron Synapses.” Science 338 (6106): 536–40.

82. Szabadics, János, Csaba Varga, Gábor Molnár, Szabolcs Oláh, Pál Barzó, and Gábor Tamás. 2006. “Excitatory Effect of GABAergic Axo-Axonic Cells in Cortical Microcircuits.” Science 311 (5758): 233– 35.

83. Szegedi, Viktor, Gábor Molnár, Melinda Paizs, Eszter Csakvari, Pál Barzó, Gábor Tamás, and Karri Lamsa. 2017. “High-Precision Fast-Spiking Basket Cell Discharges during Complex Events in the Human Neocortex.” eNeuro 4 (5). https://doi.org/10.1523/ENEURO.0260-17.2017.

84. Szegedi, Viktor, Melinda Paizs, Judith Baka, Pál Barzó, Gábor Molnár, Gabor Tamas, and Karri Lamsa. 2020. “Robust Perisomatic GABAergic Self-Innervation Inhibits Basket Cells in the Human and Mouse Supragranular Neocortex.” eLife 9 (January). https://doi.org/10.7554/eLife.51691.

85. Szegedi, Viktor, Melinda Paizs, Eszter Csakvari, Gabor Molnar, Pal Barzo, Gabor Tamas, and Karri Lamsa. 2016. “Plasticity in Single Axon Glutamatergic Connection to GABAergic Interneurons Regulates Complex Events in the Human Neocortex.” PLoS Biology 14 (11): e2000237.

86. Tasic, Bosiljka, Zizhen Yao, Lucas T. Graybuck, Kimberly A. Smith, Thuc Nghi Nguyen, Darren Bertagnolli, Jeff Goldy, et al. 2018. “Shared and Distinct Transcriptomic Cell Types across Neocortical Areas.” Nature 563 (7729): 72–78.

87. Testa-Silva, Guilherme, Matthijs B. Verhoog, Daniele Linaro, Christiaan P. J. de Kock, Johannes C. Baayen, Rhiannon M. Meredith, Chris I. De Zeeuw, Michele Giugliano, and Huibert D. Mansvelder. 2014. “High Bandwidth Synaptic Communication and Frequency Tracking in Human Neocortex.” PLoS Biology 12 (11): e1002007.

88. Tian, Cuiping, Kaiyan Wang, Wei Ke, Hui Guo, and Yousheng Shu. 2014. “Molecular Identity of Axonal Sodium Channels in Human Cortical Pyramidal Cells.” Frontiers in Cellular Neuroscience. https://doi.org/10.3389/fncel.2014.00297.

89. Ting, Jonathan T., Brian Kalmbach, Peter Chong, Rebecca de Frates, C. Dirk Keene, Ryder P. Gwinn, Charles Cobbs, et al. 2018. “A Robust Ex Vivo Experimental Platform for Molecular-Genetic Dissection of Adult Human Neocortical Cell Types and Circuits.” Scientific Reports 8 (1): 8407.

90. Toledo-Rodriguez, Maria, and Henry Markram. 2014. “Single-Cell RT-PCR, a Technique to Decipher the Electrical, Anatomical, and Genetic Determinants of Neuronal Diversity.” Methods in Molecular Biology. https://doi.org/10.1007/978-1-4939-1096-0_8.

91. Tremblay, Robin, Soohyun Lee, and Bernardo Rudy. 2016. “GABAergic Interneurons in the Neocortex: From Cellular Properties to Circuits.” Neuron 91 (2): 260–92.

92. Verhoog, Matthijs B., Natalia A. Goriounova, Joshua Obermayer, Jasper Stroeder, J. J. Johannes Hjorth, Guilherme Testa-Silva, Johannes C. Baayen, Christiaan P. J. de Kock, Rhiannon M. Meredith, and Huibert D. Mansvelder. 2013. “Mechanisms Underlying the Rules for Associative Plasticity at Adult Human Neocortical Synapses.” The Journal of Neuroscience. https://doi.org/10.1523/jneurosci.3158-13.2013.

93. Vitureira, Nathalia, Mathieu Letellier, Ian J. White, and Yukiko Goda. 2012. “Differential Control of Presynaptic Efficacy by Postsynaptic N-Cadherin and β-Catenin.” Nature Neuroscience. https://doi.org/10.1038/nn.2995.

94. Vormstein-Schneider, Douglas, Jessica D. Lin, Kenneth A. Pelkey, Ramesh Chittajallu, Baolin Guo, Mario A. Arias-Garcia, Kathryn Allaway, et al. 2020. “Viral Manipulation of Functionally Distinct Interneurons in Mice, Non-Human Primates and Humans.” Nature Neuroscience 23 (12): 1629–36.

95. Wang, Bo, Luping Yin, Xiaolong Zou, Min Ye, Yaping Liu, Ting He, Suixin Deng, et al. 2015. “A Subtype of Inhibitory Interneuron with Intrinsic Persistent Activity in Human and Monkey Neocortex.” Cell Reports 10 (9): 1450–58.

96. Wang, Yuhan, Mark Eddison, Greg Fleishman, Martin Weigert, Shengjin Xu, Tim Wang, Konrad Rokicki, et al. 2021. “EASI-FISH for Thick Tissue Defines Lateral Hypothalamus Spatio-Molecular Organization.” Cell 184 (26): 6361–77.e24.

97. Waters, Jack, and Stephen J. Smith. 2002. “Vesicle Pool Partitioning Influences Presynaptic Diversity and Weighting in Rat Hippocampal Synapses.” The Journal of Physiology 541 (Pt 3): 811–23.

98. Wit, Joris de, and Anirvan Ghosh. 2016. “Specification of Synaptic Connectivity by Cell Surface Interactions.” Nature Reviews. Neuroscience 17 (1): 22–35.

99. Wu, Tianzhi, Erqiang Hu, Shuangbin Xu, Meijun Chen, Pingfan Guo, Zehan Dai, Tingze Feng, et al. 2021. “clusterProfiler 4.0: A Universal Enrichment Tool for Interpreting Omics Data.” Innovation (New York, N.Y.) 2 (3): 100141.

100. Ximerakis, Methodios, Scott L. Lipnick, Brendan T. Innes, Sean K. Simmons, Xian Adiconis, Danielle Dionne, Brittany A. Mayweather, et al. 2019. “Single-Cell Transcriptomic Profiling of the Aging Mouse Brain.” Nature Neuroscience. https://doi.org/10.1038/s41593-019-0491-3.

101. Zeisel, Amit, Ana B. Muñoz-Manchado, Simone Codeluppi, Peter Lönnerberg, Gioele La Manno, Anna Juréus, Sueli Marques, et al. 2015. “Brain Structure. Cell Types in the Mouse Cortex and Hippocampus Revealed by Single-Cell RNA-Seq.” Science 347 (6226): 1138–42.

102. Zerucha, T., T. Stühmer, G. Hatch, B. K. Park, Q. Long, G. Yu, A. Gambarotta, J. R. Schultz, J. L. Rubenstein, and M. Ekker. 2000. “A Highly Conserved Enhancer in the Dlx5/Dlx6 Intergenic Region Is the Site of Cross-Regulatory Interactions between Dlx Genes in the Embryonic Forebrain.” The Journal of Neuroscience: The Official Journal of the Society for Neuroscience 20 (2): 709–21.

